# Seeking smarts: Male chickadees with better spatial cognition sire more extra-pair young

**DOI:** 10.1101/2025.09.30.679593

**Authors:** Carrie L Branch, Benjamin R Sonnenberg, Joseph F Welklin, Bronwyn G Butcher, Virginia K Heinen, Angela M Pitera, Lauren M Benedict, Eli S Bridge, Irby J Lovette, Michael S Webster, Vladimir V Pravosudov

## Abstract

Across animal taxa, females commonly mate with more than one male, even in monogamous mating systems. These extra-pair copulations and resulting young may increase the fitness of the female via a variety of mechanisms. The ‘good genes’ hypothesis suggests that socially monogamous females mate outside their pair bond to increase the fitness of their offspring via indirect genetic benefits, because extra-pair males do not provide parental care. We tested this hypothesis by quantifying extra-pair paternity in nonmigratory, food-caching mountain chickadees (*Poecile gambeli*). Chickadees rely on spatial cognition to recover scattered food caches and variation in spatial cognition is associated with increased survival, longer lifespan, and is heritable. However, less is known about the role of sexual selection on spatial cognitive abilities. In the current study, we found that 1. males with better spatial cognitive abilities sired more extra-pair young and produce heavier offspring in their own nests compared to their poorer performing counterparts, and 2. extra-pair males had significantly better spatial cognition than the social males they cuckolded. These results suggest that sexual selection shapes the evolution of spatial cognition in food-caching chickadees and are consistent with the good genes hypothesis, which posits that females gain indirect genetic benefits via extra-pair young.

## Introduction

The role of sexual selection and the consequences that follow from mating decisions are central focuses of evolutionary biology (1–5). Numerous hypotheses have been generated to explain patterns of mate choice, including those associated with the evolution of seemingly arbitrary attractiveness (e.g., runaway selection (6,7)) or sensory exploitation (e.g., sensory drive (8)), and those at the intersection of sexual and natural selection, when sexually selected traits have direct fitness consequences (e.g., good genes (9)). Although it has been recognized for some time that males can be actively involved in mate choice (10,11), considerable work shows that females tend to be the choosier sex, as females often invest more in gamete production and parental care (12–15). Multiple enduring hypotheses argue that females should prefer males of ‘high-quality’, frequently indicated by secondary sexual signals including ornaments and courtship displays (9, 14, 16).

Throughout the literature, male quality refers to everything from sperm quality (17) and parasite load (18), to quality of parental care, nuptial gifts, and territories (e.g., direct benefits; 19-21), to ‘good genes’ that offspring are likely to inherit (e.g., indirect benefits; 9, 16, 22, 23). Untangling these potential targets of sexual selection requires an understanding of the fitness-related traits of interest, including the direction and type of fitness consequences (e.g., survival or reproduction), as well as whether variation in the trait is genetically heritable, so that the ‘high-quality’ of a parent can be inherited by the offspring (24). Identifying and quantifying fitness-related traits in the wild that are associated with sexual selection and mate choice is challenging especially for complex behaviors, like cognition (25).

Cognitive abilities, including associative learning and memory, allow animals to succeed across variable environments, providing a buffer against unpredictable availability of resources (26, 27). If variation in cognitive abilities is associated with differential fitness outcomes (survival, reproduction, or both), and this variation is heritable, natural selection can shape cognitive variation (24). Furthermore, when heritable cognitive variation directly confers a fitness advantage, it may be associated with mate preference via secondary sexual signals or by direct observation of mate behavior (28–32). For example, females of several species show preferences for males with better cognition in lab-based, pairwise choice paradigms (30–32), while in the wild, males with better cognition have been shown to produce more young (33, 34). However, there is scant evidence from the wild that females prefer males with better cognition (28), and little evidence outside of model organisms that shows heritability of cognitive abilities (35).

In the current study, we investigated the potential role of sexual selection on the evolution of spatial cognitive abilities by examining extra-pair paternity in a wild population of socially monogamous mountain chickadees (*Poecile gambeli*). Mountain chickadees are nonmigratory, food caching birds that rely on specialized spatial cognition to recover thousands of scattered food stores throughout their territories, enabling them to survive harsh montane winters (36, 37). Following the breeding season, in the temperate autumn, juvenile chickadees are recruited into flocks of unrelated male-female pairs (36). Breeding pairs from the recent breeding season typically stay together, providing little opportunity for social mate choice at this time; however, being paired within the flock is critical because it increases overwinter survival, particularly for females (38, 39).

Work in our long-term study system using ‘smart’ feeder arrays (Figure 1) shows that individual variation in spatial learning and memory abilities confers a fitness advantage in wild chickadees: individuals with better spatial cognitive abilities are more likely to survive the winter and live longer compared to those with worse abilities (40, 41) and in resource abundant years, females lay larger clutches and fledge larger broods when their social male exhibits better spatial learning and memory abilities (42). That said, social pairs are not formed assortatively based on cognitive abilities (43), potentially due to the constraints on social mate choice mentioned above. Previous research in chickadees shows that females often produce extra-pair young with socially dominant males (e.g., 44, 45), however, in our study system we have not observed a relationship between social dominance and spatial cognitive abilities (46). One potential explanation for this, is that social dominance changes over an individual’s lifetime and with age, while spatial cognition is repeatable and consistent over time (25, 27–32).

**Figure 1.**
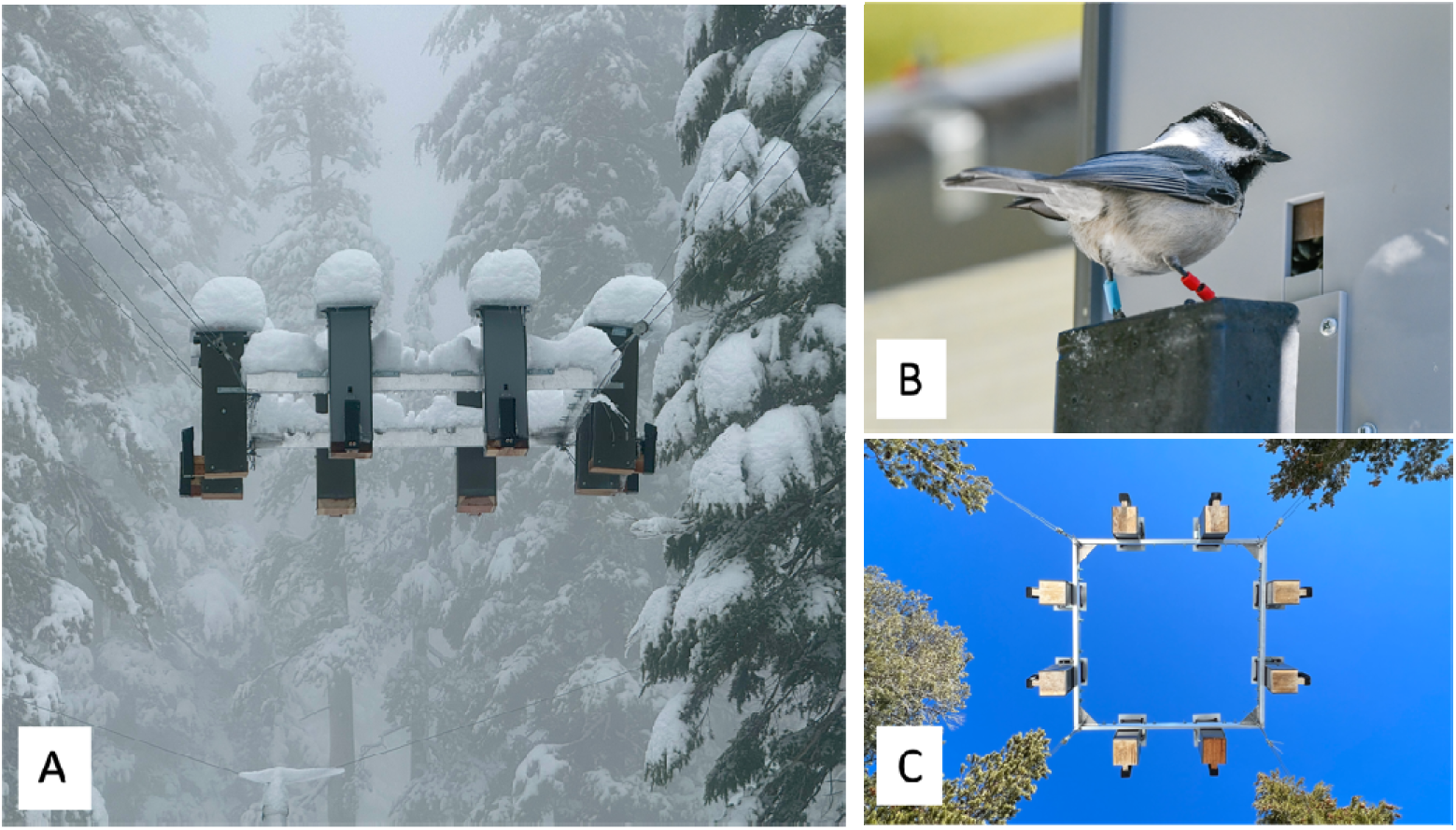
Photographs of spatial cognitive testing apparatus. A. Side-view of one array suspended in the air. B. A mountain chickadee tagged with a passive integrated transponder (PIT) tag visiting the ‘smart’ feeder array. Motorized door is open. The blue band on the left leg of the bird is the PIT tag. C. View of the testing apparatus from below, depicting equidistant placement of eight ‘smart’ feeders. Data presented come from four independent feeder arrays.

Using genome-wide association and gene network approaches, we have shown that individual variation in spatial cognitive abilities is associated with genetic differences, including genes involved in the development and function of the hippocampus, a brain region involved in spatial cognitive function (47, 48). Finally, we have shown that spatial cognitive abilities remain stable across birds’ natural lifespan (e.g., no detectable senescence (49) and does not change with age (40, 41)). Due to their life-history and reliance on specialized spatial cognition, chickadees provide a great system for studying the role of sexual selection on a cognitive trait in the wild.

Given that individual variation in spatial cognitive abilities in chickadees is genetically heritable and better cognitive abilities lead to increased overwinter survival and longevity, we predicted that spatial cognition should be an important target of sexual selection via extra-pair mating. The good genes hypothesis predicts that females should seek extra-pair copulations when their social mate is of lower quality and with a male that is of better quality than their social male (1, 9, 50, 51). We tested this hypothesis using three primary approaches: 1. Assessing the relationship between male spatial cognition and the number of extra-pair young sired, 2. Assessing whether extra-pair young occurred more often in nests of males with poor spatial cognition and 3. Comparing the spatial cognition of males siring extra-pair young in a nest to the social male they cuckolded (e.g., 44, 45, 50, 52, 53). If females obtain genetic benefits via extra-pair young, we expect extra-pair males to have significantly better spatial cognition compared to the within-pair males they cuckold. In addition to testing these primary predictions, we were able to assess potential alternative factors that might explain differences in rates and directional patterns of extra-pair paternity, including effects of a. elevation, b. age, c. reproductive output, d. male parental investment, and e. female’s cognitive abilities. Finally, we mapped the location of nests based on paternity status (extra-pair vs. social) and spatial cognition to account for potential bias in male spatial distribution.

## Methods

### Experimental Design

We studied spatial learning and memory abilities and extra-pair paternity rates in mountain chickadees (*Poecile gambeli*) at our long-term field site in the northern Sierra Nevada mountains of North America (Sagehen Experimental Forest in northern California, USA, Sagehen Creek Field Station, University of California Berkeley). Since 2013, we have monitored breeding phenology in uniquely color-banded individuals across two elevations (54, 55) and measured the spatial cognitive abilities of hundreds of individuals starting in 2015 (e.g., 40-43, 46-49, 56-59). The breeding season begins in April and continues through July, while the non-breeding season spans from August to March of the following year. Mountain chickadees are secondary cavity nesting birds that readily use nest boxes at our field site. Every year in late April we begin monitoring 350 nest boxes throughout our field site for evidence of nest building. Once nesting material is detected we monitor all stages of breeding through fledging of each nest, recording information about the date the first egg is laid, start of incubation, and hatching, and record the clutch size (number of eggs), and the brood size (number of chicks at post-hatch day 16) of each nest. We process all chicks in the nest box at post-hatch day 16, which includes banding them with an aluminum United States Geological Survey (USGS) band, weighing them, and obtaining a small blood sample for genetic and paternity analyses. We trap adult and first-year birds using mist nets at established feeders during annual autumn trapping and by hand at nest boxes during the breeding season. Upon capture, birds are fitted with color-bands and a unique Passive Integrated Transponder (PIT) tag, wing length is measured, and age is estimated using multiple plumage features (60). When nestlings banded with USGS bands are recruited into our population from nest boxes, we band them with an additional color band and PIT tag for identification.

### Testing spatial cognition in the wild

Spatial learning and memory testing was conducted using two feeder arrays at each of our elevation sites (High, range: 2380 m – 2590 m; coordinates: 39.42402, -120.315015 and Low, range: 1965 m – 2070 m; coordinates: 39.443500, -120.243248), following 40-43, 46-49, 56-59. Within an elevation, the feeder arrays were established ca. 1.5 km apart with mostly non-overlapping birds visiting each array. Each array consisted of eight equidistant RFID-based feeders on a square (122 x 122 cm) aluminum frame suspended ca. 4 m above the ground (Figure 1). Each feeder is equipped with a small, motorized door. Prior to testing, we kept the feeder doors of the arrays open for approximately one month. Next, the feeder doors were closed and programmed to open for any bird with a PIT tag, this allowed birds to habituate to the motorized feeder doors for one week. Finally, we measured the spatial learning and memory abilities of mountain chickadees using a 4-day task. The spatial learning and memory task involved pseudo-randomly assigning each bird to a single feeder in the 8-feeder array that was not its most-visited feeder during the previous two habituation stages. During this task, each bird can only obtain a food reward (black oil sunflower seed) from its single assigned feeder, but all eight feeders recorded the date, time, and identity of every visit by all PIT tagged individuals. We measured cognitive performance by estimating the number of location errors (e.g., non-rewarding feeders visited prior to visiting the correct rewarding feeder, max 7) per trial, over multiple trials. A trial began when a bird visited any feeder in the array and ended with a visit to the rewarding feeder. If the bird landed on its rewarding feeder first, it was considered to have made zero location errors. Chickadees collect a single sunflower seed during each trial and fly away from the array to nearby trees to either eat or cache the seed before returning to the array and each trial takes just a few seconds (58).

We used the mean number of location errors per trial over the first 20 trials and the mean number of location errors per trial over the entire 4-day task as the two metrics of cognitive performance, following our previous work (40–43, 46–49, 56–59). These measurements capture learning curves in a single metric (Figure S7); as birds learn the rewarding feeder location, they make fewer and fewer errors on each consecutive trial. Using the mean number of location errors over multiple trials captures this improvement – the mean number of location errors per trial over multiple trials is smaller for individuals that improved faster and larger for individuals that improved slower. While these metrics are correlated (Figure S7), the relationship is not 1 to 1. Performance on the first 20 trials of the task reflects learning acquisition, while performance on the entire task reflects memory retention (40). Since birds completed different numbers of trials over the entire task, we used the total number of trials as a covariate in models involving performance on the entire 4-day task (40–43, 46–49, 56–59). We have previously documented that variation in both metrics is associated with significant differences in survival (40), longevity (41), in female reproductive decisions (42), in foraging decisions (58), and in cognitive flexibility (59). For analyses of spatial cognitive abilities and extra-pair paternity we used cognitive performance measured in the first year each individual completed the task, as cognitive performance on the spatial learning and memory task does not change with age (40, 57). In addition, birds were included in analyses if they performed at least 20 trials and made fewer than mean 3 errors across the 4-day testing period. Males that bred in more than one year were included in the analyses (controlled for using male ID as a random effect); as such when reporting results, we report the sample size for unique nests and ‘male-years’.

### Assessing extra-pair paternity

#### DNA extraction and ddRAD sequencing

For this study, we aimed to sequence and genotype parents with cognition data and their nestlings over four consecutive years (2018–2021). Each year, we prioritized sequencing nests of males and potential extra-pair sires that performed in our cognitive task. Samples from each year were sequenced at the end of each breeding season. Unfortunately, genotypes from 2019 were of low quality and paternity assignments could not be completed with confidence, therefore we present data from 2018, 2020, and 2021.

Blood samples from adults and nestlings were stored at room temperature in Queens Lysis buffer until DNA was extracted using Qiagen DNeasy Blood & Tissue kits. The DNA concentration of each sample was quantified using Qubit Fluorometer 2.0 (Life Technologies) prior to library preparation and sequencing. We used a double digest restriction-site associated DNA (ddRAD) sequencing approach from (61) to obtain single nucleotide polymorphisms (SNPs) for our paternity analyses. Sequencing was completed in batches following field sampling of each year, therefore, individuals from each breeding year were run together, but separately from the other years. For each individual, 100-500 ng of extracted DNA was digested with restriction enzymes SbfI-HF (NEB) and MspI (NEB) and ligated with one of 20 unique P1 adapters and a P2 adapter (P2-MspI). Next, samples were pooled by groups of 20, maintaining individual identity via unique P1 adapters. Samples were then purified using 1.5x volume of homemade solid-phase reversible immobilization (SPRI) beads, made with Sera-Mag SpeedBead Carboxylate-Modified Magnetic Particles (Cytiva) (62). These pooled samples (index groups) were sent to Cornell University Biotechnology Resource Center (BRC; RRID:SCR_021727) for size selection between 450 – 600 bp using BluePippin (Sage Science). Following size selection, adapter ligated DNA was enriched by 11 cycles of polymerase chain reaction (PCR) with a universal i5 primer and a unique indexed i7 primer for each index group using Phusion DNA polymerase (NEB). Reactions were again cleaned using 0.7x volume of homemade SPRI beads (as above) and the index groups pooled in equimolar ratios to create a single sequencing library run on one lane of Ilumnia NextSeq500 (150 bp single end, performed at BRC). Sequencing was performed with a ∼20% PhiX spike-in to introduce diversity into the library.

#### SNP Analysis and Paternity assignment

##### Quality filtering, demultiplexing, and genome alignment

The following were run the same, but separately for each year of samples. First, the quality of all reads were assessed using FASTQC version 0.11.9 (https://www.bioinformatics.babraham.ac.uk/projects/fastqc/). All sequences were trimmed using *fastx_trimmer* (FASTX-Toolkit) to a length of 147 bp on the 3’ end to exclude low-quality SNP calls. Next, reads containing a single base with a Phred quality score less than 10 were removed and any sequences that had more than 5% of the bases with a Phred quality score lower than 20 were also removed. We then used *process_radtags* from STACKS version 2.65 to demultiplex the sequence reads and obtain files with specific sequences for each individual. Next, we aligned sequences to the chromosome scale reference genome of mountain chickadees (48) using Bowtie2 version 2.5.1. The resulting .sam files were converted to .bam files and sorted using the *view* and *sort* functions in Samtools version 1.18. These aligned reads were used as the input for the *ref_map.pl* program in Stacks 2.65.

##### SNP filtering and assigning paternity

SNPs were exported using *populations* in Stacks 2 version 2.65, and each year’s samples were run separately to maximize the number of SNPs and individuals included for downstream paternity analyses. All male individuals alive during the year of interest were included as potential sires. Individuals with more than 50% missing SNP data were removed from the dataset; only 5 individuals from the 2021 dataset did not meet this criterion. Loci were exported if they were present in 85% of that year’s sample population (–*r*). If a RAD locus had more than one SNP, data were restricted to the first SNP *(–write_single_snp*) to minimize those in high linkage disequilibrium. We required a minor allele frequency of 0.05 to process a nucleotide *(– min_maf),* then filtered for loci in Hardy-Weinberg equilibrium (p<0.05) using VCFtools (*--hwe* option) (63,64). Variant call format (VCF) files were converted to GENEPOP format using the ‘radiator’ package in R (65).

Before assigning paternity, we imported the yearly GENEPOP files into CERVUS (66), then ran maternity tests separately for each elevation in each year to confirm mother-offspring relationships identified during field observations. We assessed the quality of these assignments by plotting the distributions of the first and second-ranked mother’s scores for each offspring. First and second-ranked mother pair LOD scores formed bimodal distributions, with nearly all first-ranked mother scores above zero and nearly all second-ranked mother scores below zero. In rare cases when second-ranked mother scores were positive, these were almost always known sisters or daughters of the mother identified during field observations. Therefore, first-ranked mother-offspring pairs with positive log likelihood (LOD) scores were considered correct assignments, but pairs with negative assignments were not included in paternity analyses because we were not confident in the mother assignment. Offspring resulting from a female laying an egg in a nest that was not her own (i.e., egg dumps) were uncommon and not included in further statistical analyses.

Paternity analyses were conducted in CERVUS separately for each elevation in each year by calculating trio LOD scores for each potential father-offspring pair. Trio LOD scores account for known maternal genotypes when identifying potential fathers. As we did for mother-offspring pairs, we plotted the distribution of the first and second-ranked male scores and again found a bimodal distribution centered around zero (Figure S8). Based on these plots, we set cut-off scores to assign paternity by: all first-ranked male scores at an elevation in a year that were positive and greater than the top second-ranked male score at that elevation in that year were assigned as fathers. Offspring sired by the social male at the nest were considered within-pair, whereas offspring sired by other males were considered extra-pair. Offspring whose social father was genotyped, but we could not assign a father to, were also considered extra-pair.

##### Statistical analysis

We tested whether the mean number of location errors per trial across the first 20 trials correlated with the mean number of location errors per trial over the entire 4-day task for males using a linear model. Mean location errors per trial over the entire task was used as the response variable while mean location errors over the first 20 trials and the total number of trials completed were included as fixed effects. Numbers and letters of subheadings correspond to predictions in last paragraph of the Introduction and carry throughout the Results.

###### 1. Testing effect of male spatial cognition on the number of extra-pair young sired

We employed two methods to test whether spatial cognitive ability was associated with the number of extra-pair young a male sired in a year. Both models included the number of extra-pair young sired as the response variable, with spatial cognitive performance (refers to separate models run for (1) the mean number of location errors per trial over the first 20 trials (first 20) and (2) the mean number of location errors per trial over the 4-day task (entire task)), elevation, and year as fixed effects, and male identity as a random effect to control for repeated measures of males across years. Both models used the Poisson error distribution and a ‘log’ link, but each model accounted for zeros in the dataset differently. The first method included a zero-inflation parameter (ziformula∼1), using the R package ‘glmmTMB’ (67) to account for observations of males with zero extra-pair young, whereas the second method restricted the dataset to males that sired at least one extra-pair offspring.

###### a. Effect of elevation on number of extra-pair young sired

Much of the work in our mountain chickadee system has focused on elevation-related differences in spatial learning and memory performance, therefore, we tested whether extra-pair paternity rates differed between the high and low elevations using generalized mixed effects models with binomial error distributions and ‘logit’ links, using the R package ‘glmmTMB’ (67). Our response variable was whether each offspring was sired by an extra-pair father or not and we only included nests in which more than 50% of the offspring were genotyped and run through the paternity analysis. Elevation was included as a fixed effect and mother identity was included as a random effect in each model. A likelihood ratio test showed statistical support for an interaction between elevation and year (p<0.05), so we tested the effect of elevation on extra-pair paternity rates separately for each year.

###### b. Effect of male age on number of extra-pair young sired

It is possible that a relationship between extra-pair paternity and spatial cognitive ability is driven by age, as older males often sire more extra-pair offspring than younger males in many systems (e.g., 68). While chickadees with better spatial cognition in our study system tend to live longer, cognitive performance does not change with age in the same individuals (40, 41, 57). However, we tested whether including male age as a fixed effect improved each model using likelihood ratio tests that compared models with and without male age.

###### c. Effects of spatial cognition on brood size and nestling mass

In addition, cognitive ability could come at a cost to breeding performance, so we tested whether spatial cognitive ability was associated with breeding performance using a generalized linear mixed model with the generalized Poisson distribution (69). Brood size was included as the response variable and male and female spatial performance on the first 20 trials, elevation, and clutch size were included as fixed effects. For these analyses, we used all breeding data available, therefore, year was included as a random effect in this analysis because it had enough levels (2015–2023). Male and female identity were also included as random effects to control for yearly variation and repeated measures of individual males and females. In a second analysis, we used a linear model that included mean nestling mass at day 16 as the response variable (in our system, mass of nestlings on day 16 is a significant predictor of survival and recruitment (70)), and included male and female performance on the first 20 trials of the spatial cognitive task and elevation as fixed effects, and year (2015–2023) and male and female identity as random effects. Again, year (2015–2023) and male and female identity were included as random effects to control for yearly variation and repeated measures of individual males and females.

###### d. Effect of paternity status on male investment

Siring extra-pair offspring may reduce a male’s investment in his own nest, so we tested whether siring extra-pair young was associated with breeding performance. We used a generalized linear mixed model with the generalized Poisson distribution to test whether brood size in a male’s nest was affected by the male siring extra-pair offspring (yes or no), and we used a linear mixed model to test whether the mean mass of the nestlings at day 16 in a male’s nest was affected by whether the male sired any extra-pair offspring (using three years of data on extra-pair paternity). Elevation, clutch size, and year (three years of extra-pair paternity data) were included as fixed effects and male and female identity were included as random effects in both models to control for repeated measures of individual males and females. We tested whether the mass of young sired by extra-pair males differed from the mass of young sired by social males using a linear mixed model with nestling mass at day 16 as the response variable and extra-pair status, elevation, and year as fixed effects. Nest identity was included as a random effect to control for nestlings that came from the same nests.

###### 2. Testing effect of male spatial cognition on likelihood of being cuckolded

We also tested whether males with worse spatial cognitive abilities were more likely to be cuckolded than those that performed better on the task. Male spatial cognition was included as the response variable and elevation and whether the nest had any extra-pair young were included as fixed effects. When modeling the mean number of location errors per trial on the entire task we used the Gamma distribution and included the total number of trials completed as a fixed effect. Including male identity as a random effect in both models led to convergence errors, so male identity was not included in the models.

###### e. Effect of female cognition on number of extra-pair young

Both male and female cognitive abilities could affect the likelihood of a nest containing extra-pair young, so we tested whether cuckoldry was associated with the spatial cognitive abilities of both parents at a nest using multiple methods. First, we modeled the relationship between male and female spatial cognitive ability and cuckoldry at the nest and offspring level. At the nest level, we tested whether male or female spatial cognitive ability predicted whether any offspring in a nest were sired by extra-pair males. At the offspring level, we tested whether male or female spatial cognitive ability predicted whether each offspring was sired by an extra-pair male or not. For both tests, we used generalized linear mixed models with binomial distributions that included male and female spatial cognitive performance, elevation, and year as fixed effects. Male and female identity were included as random effects to control for repeated measures of individual males and females.

Second, for nests that had at least one extra-pair young, we tested if the number of extra-pair young in a nest was associated with male or female spatial cognitive ability using a generalized linear mixed model with a truncated Poisson distribution. The number of extra-pair young in the nest was included as the response variable, while male and female performance on the first 20 trials, year, and elevation were included as fixed effects. Male and female identity were included as random effects to control for repeated measures of individual males and females. The effect of male age on cuckoldry rates was tested by adding male age as a fixed effect to each model and comparing models with and without male age using likelihood ratio tests.

###### 3. Comparing spatial cognition of extra-pair males and the social males they cuckold

If males initiate extra-pair copulations, we expect them to be randomly distributed across different nests, since the fitness benefit of extra-pair young is increased for the male regardless of the cognitive abilities of the females’ social mate. On the other hand, if females initiate extra-pair copulations, we expect them to select males of better quality than their social mate – in our case, males with better spatial cognition – as this could result in indirect benefits to the female via offspring with better cognition and higher likelihood of survival (27, 28). Therefore, we tested whether the spatial cognitive abilities of extra-pair males differed from those of the males they cuckolded by comparing spatial cognition between father types (extra-pair male vs social male within the same nest) using linear mixed models. Spatial cognitive performance was used as the response variable, then father type (extra-pair mate versus social mate) and elevation were included as fixed effects. Offspring identity was included as a random effect to restrict comparisons to within offspring and mother identity was included as a random effect to control for repeated measures of females. When fitting models to the total location errors, we log-transformed the response variable to improve residual fit and included a quadratic term of total trials completed during the 4-day cognitive testing period, since the number of trials completed influences performance (35, 40, 41, 57).

###### b. Comparing age of social and extra-pair males

Finally, we tested whether extra-pair males were significantly older than the social males they cuckolded using a chi-square test and paired t-test.

All statistical analyses were performed in R version 4.1.1 (71) and the residual fit of all models was checked using DHARMa (72). Statistical significance was calculated using Wald Chi-square tests.

## Results

There was a strong positive correlation between the number of location errors on the entire 4-day task (entire task) and on the first 20 trials (first 20) for males completing the task in our study (Coef=0.27, F=121.72, P<0.001; Figure S7a).

### Extra-pair paternity and spatial cognition

Across three years of data collection (2018, 2020, and 2021), we were able to genotype and assign paternity to 86% of offspring in the study (630 of 732). We found that 30.3% of the total offspring sampled (222 of 732) were sired by extra-pair males and that 70.1% of nests (89 of 127) had at least one extra-pair young (extra-pair young). Numbers and letters of subheadings correspond to predictions in last paragraph of the Introduction and Statistical Methods section.

#### 1. Males with better spatial cognition produced more extra-pair young

Performance on the spatial learning and memory task predicted the number of extra-pair young a male sired each year (Figure 2 and Table S3). Males with better spatial learning and memory abilities (e.g., fewer mean location errors per trial) sired more extra-pair young, both according to a zero-inflated model including males that sired no extra-pair young (first 20 trials: ^2^ =6.28, P=0.012 (Figure 2, dashed line); entire task: χ^2^_1_=8.12, P=0.004, N=280 (137 individual males; Figure S5)), and to a model only including males that sired at least one extra-pair young (first 20 trials: χ^2^_1_=5.96, P=0.015 (Figure 2, solid line); entire task: χ^2^_1_=7.68, P=0.006, N=41 (34 individual males; Figure S5). This model suggests that of the males that sired at least one extra-pair young, those with the best cognitive performance may sire between 6 and 7 extra-pair young each year, whereas males with the worst cognitive performance will sire only 1 to 2 extra-pair young each year. We have shown that birds with better spatial cognitive abilities live longer (41) and lack reproductive senescence within their natural life span (73), therefore, males with better cognition are expected to produce significantly more offspring over their lifetime.

**Figure 2.**
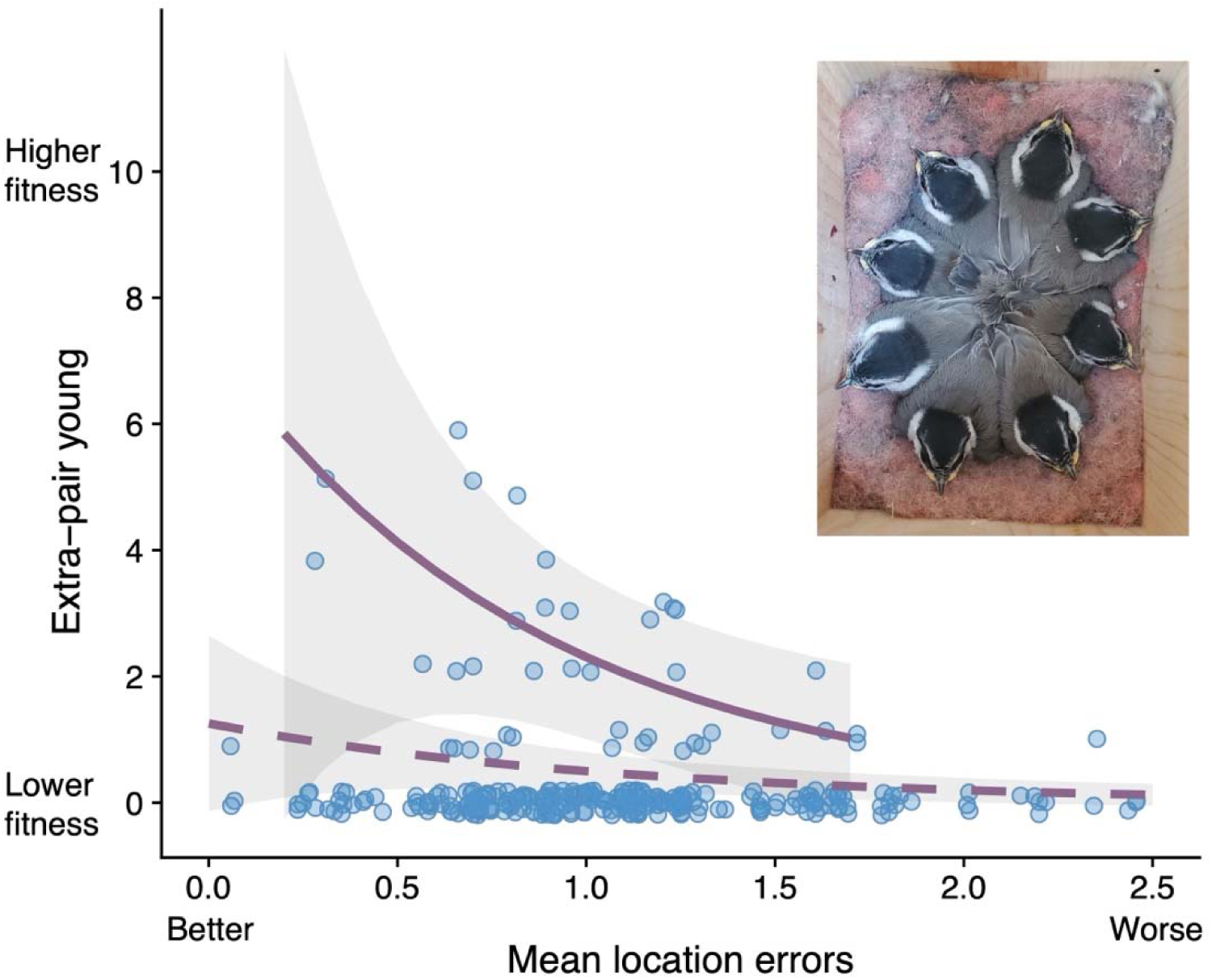
Model predictions of the number of extra-pair young a male sired in a year by the mean number of location errors per trial over the first 20 trials. The dashed line represents a zero-inflated model for all males, including those with no extra-pair young (N=280; 137 individual males), and the solid line represents the model prediction for males that sired at least one extra-pair young (N=41; 34 individual males). Points are slightly *jittered* on the y-axis to improve readability. Shading represents bootstrapped 95% confidence intervals. Inset depicts 8 nestlings at 16 days post-hatching. See model predictions for the number of extra-pair young a male sired in a year by total 4-day testing performance in Figure S5.

##### a. Effect of elevation

We found no significant differences in the number of extra-pair young sired at high versus low elevation populations regardless of year sampled (Supplemental Results, Figure S1, Table S1 and S2), therefore, we combined high and low elevations for all later analyses. Data from breeding birds in this study confirmed previous research showing that female chickadees seek extra-pair copulations with neighboring males (on average ∼100 m from their own nests; Figures S2 and S3), which are likely familiar males from their nonbreeding flocks (36, 39, 44, 74). In addition, breeding male territories do not appear to cluster based on cognitive performance (Figure S4).

##### b. Effect of age

Importantly, we show that male age was not associated with the number of extra-pair young sired (see Supplemental Results).

##### c. Effects of reproductive output

In addition to testing how cognitive abilities were related to number of extra-pair young, we also tested whether variation in cognitive abilities had direct consequences on breeding performance. Across 9 years of breeding data (2015–2023), we detected no relationship between the social males’ spatial cognitive abilities and brood size (χ^2^_1_=2.32, P=0.127, N=205 nests), or the females’ spatial cognitive abilities and brood size (χ^2^_1_=1.75, P=0.186, N=205 nests). Controlling for the effect of clutch size on brood size produced similar results (Table S4). However, males with better performance on the first 20 trials raised heavier young (mass at day 16, χ^2^_1_=4.86, P=0.028, N=202 nests, Table S4 for entire task), while female performance on the first 20 trials showed no significant relationship with offspring mass at day 16 (χ^2^_1_=0.93, P=0.335, N=202 nests, Table S4 for entire task).

##### d. Effect of male investment

Three years of extra-pair paternity data suggest that extra-pair males do not compromise their own nest investment compared to males that did not sire any extra-pair offspring, as neither brood size (brood size: χ^2^_1_=0.30, P=0.584, N=144 nests) nor mean nestling mass (mean mass: χ^2^_1_=2.10, P=0.147, N=140 nests) differed significantly between males that had extra-pair young and males that did not sire any extra-pair young (Table S5). We also compared the mass of young sired by extra-pair males to those sired by the social males raised in the same nests and found no significant difference ( χ^2^_1_=2.83, P=0.093, N=719 offspring, Table S6).

#### 2. Does spatial cognition of the social male predict presence of extra-pair young?

Across three years of data collection, we had spatial cognition data on the social males of 101 nests; approximately 28% of nests (28/101) had no extra-pair young and 72% of nests had at least one extra-pair young (73/101). The spatial cognitive abilities of males that had extra-pair young in their nests did not differ from males that had no extra-pair young in their nests (first 20 trials: χ^2^_1_=0.42, P=0.516; entire task: χ^2^_1_=2.07, P=0.150, N=101 nests, Table S8). Among nests where both the male and female had spatial cognition data, the likelihood of being cuckolded was not significantly related to the cognitive abilities of the social males (Offspring level: first 20: χ^2^_1_=0.25, P=0.620, entire task: χ^2^_1_=0.43, P=0.513, N=348 offspring for 42 males; Nest level: first 20: χ^2^_1_=0.05, P=0.808; entire task: χ^2^_1_=0.22, P=0.642, N=59 nests for 42 males, Table S8).

Similarly, for nests that had at least one extra-pair young, the cognitive abilities of the social male did not predict the number of extra-pair young in their nests (first 20: χ^2^_1_=1.03, P=0.311; entire task: χ^2^_1_=0.40, P=0.528, N=41 nests for 35 males, Table S9).

##### b. Effect of age

In addition, male age did not significantly affect the likelihood of a male being cuckolded (see Supplemental Results).

##### e. Effect of female cognition on extra-pair young

A female’s cognitive abilities may influence her decision to seek extra-pair copulations (i.e., a female may seek extra-pair copulation to compensate for her own poor cognition). Therefore, we assessed whether the likelihood of having extra-pair young in the nest was related to the spatial cognitive abilities of the female. Females who performed worse on the first 20 trials of the spatial cognitive task were more likely to have extra-pair young in their nests (Offspring level: first 20: χ^2^_1_=6.21, P=0.013, N=348 offspring for 42 females), though this relationship was not significant when we analyzed performance across the entire 4-day task (χ^2^_1_=0.18, P=0.673, N=348 offspring for 42 females), or for either metric at the nest level of analysis (first 20: χ^2^_1_=0.21, P=0.645, entire task: χ^2^_1_=0.13, P=0.723, N=59 nests for 42 females, Table S8).

Similarly, for nests that had at least one extra-pair young, females that performed worse on the first 20 trials had significantly more extra-pair young in their nests (first 20: χ^2^_1_=6.79, P=0.009, N=41 nests for 32 females), but there was no significant relationship across the entire 4-day task (entire task: χ^2^_1_=1.04, P=0.308, N=41 nests for 32 females, Table S9).

#### 3. Extra-pair males have better spatial cognition than the social males they cuckold

When there were extra-pair young in a nest, they were sired by males with better spatial learning and memory abilities (on average) compared to those of the social male they cuckolded (first 20 trials: χ^2^_1_=14.81, P<0.001, N=71 comparisons of a social male and the extra-pair male sire (Figure 3); entire task: χ^2^_1_=6.08, P=0.014, N=71 comparisons (Figure S6 and Table S7)).

**Figure 3.**
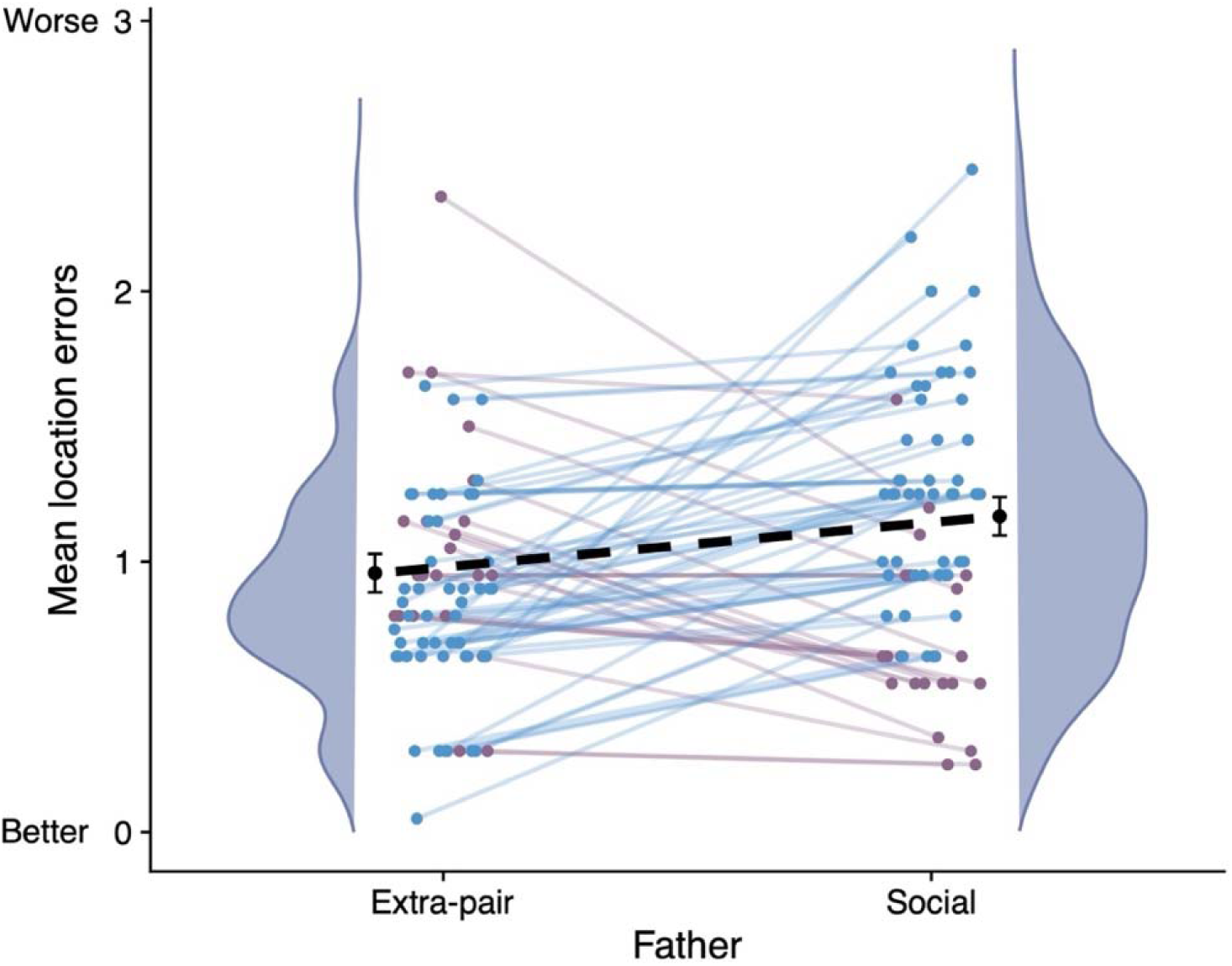
Comparing spatial cognitive performance (mean number of location errors per trial over the first 20 trials) of extra-pair (mean ± SE = 0.96 ± 0.07) and social fathers (mean ± SE = 1.17 ± 0.07, 71 comparisons). Lines connect males by offspring: blue lines show comparisons where the extra-pair father performed better than the social father (49 of 71 cases or 69%), and purple lines show comparisons where the social father performed better than the extra-pair father (19 of 71 cases, or 26.7%; in 3 cases the extra-pair and social father had the same cognition score). Shaded distributions represent smoothed histograms of cognitive performance for each father type. The same analysis is presented for performance over the 4-day testing period in Figure S6.

##### b. Effect of age

Furthermore, extra-pair males did not differ significantly in age from the males they cuckolded (28 offspring were sired by an extra-pair male that was older than the social male, 19 offspring were sired by an extra-pair male that was the same age as the social male, and 24 offspring were sired by an extra-pair male that was younger than the social male; Chi-square test: χ^2^_1_= 1.72, P=0.424). A paired samples t-test of social and extra-pair father ages returned a similar result (χ^2^_1_=0.27, P=0.662).

## Discussion

Our findings show that sexual selection acts on the spatial cognitive abilities of free-living, food-caching mountain chickadees via extra-pair paternity. We found that (a) males, but not females with better spatial cognitive abilities produce higher quality offspring (higher mass, 70) in their social nests, (b) males with better spatial learning and memory abilities sire more extra-pair young than their poorer performing counterparts and, (c) extra-pair males have better spatial cognitive abilities than the social males they cuckold, regardless of age. Importantly, our data show that absolute male cognitive abilities do not predict extra-pair young in the nest, rather that the differences in cognition between the extra-pair male and the social male are relative to each other –a good quality male can lose paternity to an even better-quality male.

We specifically quantified extra-pair paternity in our mountain chickadee system and do not have data on all extra-pair copulation attempts by males or females. However, given the extensive research on chickadee breeding behavior and their closely related Eurasian relatives (tits), we suggest these patterns of extra-pair paternity may be driven by females for several reasons; 1. Female parids (chickadees and tits) are known to pursue extra-pair copulations with neighboring males (44, 45, 74–77), 2. Parids have never been reported or observed engaging in forced copulation (e.g., 45, 74, 77), and 3. Resident males are highly territorial and do not tolerate male conspecifics in their territories. In addition, the directional relationship between the cognitive phenotype of the social and extra-pair male would not be predicted by male choice but is consistent with good genes hypotheses, where females seek high quality males (in this case, those with better spatial cognitive abilities) for extra-pair copulation (1, 9, 14). In other words, males would benefit from siring extra-pair young with any female regardless of her social male’s cognitive abilities.

That said, active choice need not be implied, as males with poor spatial cognitive abilities could be worse at defending their territories from other males, which may allow those males with better cognitive abilities to intrude and copulate with the resident female. We think this explanation is unlikely as males with worse cognitive abilities tend to be more aggressive (78, 79) and cognitive performance is not associated with social dominance (46). Females, on the other hand, would benefit from obtaining extra-pair copulations with males that have better spatial cognition than their social male, as this trait is heritable (47, 48) and would increase the likelihood of producing offspring with better spatial cognition, leading to higher overwinter survival (40) and longer lifespan (41). Indeed, females with poorer spatial memory acquisition (made more errors on the first-20 trials) had more extra-pair young in their nests, while social male cognitive performance did not predict extra-pair young in the nest. This suggests that females may compensate for their poor cognitive abilities via extra-pair young. Considering that alleles associated with better spatial cognitive abilities are not sex linked (47), producing extra-pair young with better quality males increases selection pressure on spatial cognitive abilities in both males and females, as both sexes rely on spatial cognition for survival (80).

Although it remains unknown whether chickadees can evaluate cognitive abilities directly or indirectly via some correlated trait(s), the life history and breeding ecology of mountain chickadees provide several possibilities. For example, female chickadees may use secondary sexual signals, such as song and plumage variation associated with spatial cognitive abilities, to identify high-performing males (1, 9, 28, 81). Recently, we have shown that males that perform better on our spatial learning and memory task sing more throughout the day (82) and exhibit brighter black eyelines (83) compared to their poorer performing counterparts. However, this variation in correlative and we have not shown that females use this information in discriminations. Alternatively, it is possible that females can directly assess a male’s ability to successfully recover food stores over winter and base mating decisions on this information in the spring (30, 38, 39). Chickadees are nonmigratory, sedentary birds that aggregate in flocks of unrelated male-female pairs throughout the nonbreeding season and break apart once breeding commences (36, 39). Data from this study confirm previous research showing that female chickadees seek extra-pair copulations with neighboring males (44, 74, 84), which would likely be males from their winter flock (36, 39). Regardless of if and how females evaluate male cognition, mating with males with better cognitive performance should increase female fitness by increasing the likelihood of offspring inheriting the specific alleles associated with better cognition, which would be expected to lead to higher offspring survival and longer lifespan (40, 41).

An alternative hypothesis for our findings is that females seek extra-pair copulations with other males as insurance against the social male’s infertility (84), again gaining indirect genetic benefits. This hypothesis does not explain the direction of our findings, because male cognitive abilities did not predict the number of offspring a male produced in his own nest (i.e., cuckolded social males still fathered chicks in their nests), suggesting that males with poorer cognition do not suffer from infertility. That said, we do not have data on the sperm quality of these birds and therefore cannot exclude differences in fertility or sperm competition between males with better and worse spatial cognition. Related, we have not detected fitness trade-offs or negative effects associated with better cognitive abilities in our system; we do not see cognitive or reproductive senescence within birds’ naturally occurring life span (49, 73). In fact, chickadees with better spatial cognition live longer allowing them to produce more offspring over their lifetime compared to chickadees with worse cognition (41).

Interestingly, a previous study in our system showed that females mated to social males with better spatial abilities lay larger clutches and fledge larger broods in years of resource abundance (42). Together with the present findings, we suggest that female mating decisions, including extra-pair copulations and reproductive investment, may vary opportunistically. When resources are abundant, females lay larger clutches if their social male has better cognitive abilities (42), and when males with better spatial cognition than her social mate are available for extra-pair copulations, they pursue those opportunities. Indeed, we found that females with poorer spatial cognition had more extra-pair young in their nests, suggesting female mating decisions could be based on some physiological signal associated with poorer cognition that serves as a compensatory mechanism. At the ultimate level, females with poorer cognition may benefit disproportionately by mating with the highest quality male they can find, which in the case of food-caching birds, likely means those with better spatial cognitive abilities (40, 41).

An additional alternative explanation for our findings is that birds with better spatial cognitive abilities have access to better territories and therefore are clustered based on cognitive abilities (85). Then, because females have extra-pair copulations with neighboring males (44, 74, 84), those neighbors would be males with better spatial cognition. However, in our study system, males do not appear to cluster based on cognitive performance (Figure S4), suggesting that females are not finding males with better cognitive abilities than their social males by chance. Furthermore, while higher social dominance status is associated with access to better territories in chickadees (86), we see no association with dominance status and spatial cognitive abilities in our system (46) or closely related black-capped chickadees (81). While it is possible that our results reflect some unmeasured trait(s) linked to spatial cognitive ability, such as boldness, aggression, or increased exploratory behavior (e.g., 32), previous data from our system do not support these additional alternative hypotheses. Chickadees with enhanced spatial cognitive abilities are not more aggressive (78) nor do they show higher rates of exploratory behavior (e.g., higher space use) (79).

Given the fitness advantages of superior spatial cognitive abilities for food-caching chickadees, how variation is maintained and why enhanced spatial cognitive abilities have not evolved to fixation may seem puzzling. While food-caching birds exhibit superior spatial cognitive abilities compared to their non-caching counterparts (87), considerable variation persists within species (56). Research from our system demonstrates the highly polygenic nature of the spatial cognitive phenotype, meaning there are multiple genetic pathways that may produce the same cognitive phenotype (47, 48, 88). In addition, selection strength varies with annual variation in winter climate, which in milder years, may allow individuals to survive and reproduce despite comparatively poor spatial cognitive abilities (40, 54–56). Finally, we have documented elevation-related differences in spatial cognition and the hippocampus of mountain chickadees (89), but there is gene flow among elevations (90) which moves allelic variation associated with poorer cognition within the population.

Overall, our study provides novel results showing that sexual selection contributes to the evolution of spatial cognitive abilities in a food-caching species, which rely on such spatial cognition for survival. We show that males with better spatial cognition exhibit higher reproductive output, via siring more extra-pair young and fledging heavier chicks, which are more likely to survive and be recruited into the breeding population (70). We also show that when there are extra-pair young in the nest, they are sired by males with better spatial cognition than the social male (on average), which suggests that females may be selectively obtaining extra-pair copulation with higher quality males; though we do not have direct experimental evidence of female choice. Whether these patterns are driven by females or males, males with better spatial cognitive abilities exhibit higher fitness by producing more offspring, adding to the evidence for selection on spatial cognitive abilities, including differential survival and longevity in food-caching birds (40, 41).

## Acknowledgements

Thank you to Qwahn Kent, Danielle Schaening-Lopez, Alyssa Nowicki, and Beck Kerdman for assistance with DNA extractions and the Cornell University BRC Bioinformatics Core Facility (RRID:SCR_021757) for providing computing power for all bioinformatics. All procedures were in accordance with University of Nevada Reno IACUC (protocols 20-11-1103-1, 20-06-1014, 20-08-1062) and California State Fish and Wildlife Scientific Collecting Permit (S-193630001-20007-001).

## Author contributions

Conceptualization: CLB, VVP

Methodology: CLB, BRS, JFW, BGB, VKH, AMP, LMB, ESB, MSW, IJL, VVP

Investigation: CLB, BRS, VKH, AMP, LMB, VVP

Visualization: CLB, JFW, VVP

Funding acquisition: CLB, VVP

Writing – original draft: CLB

Writing – review & editing: CLB, BRS, JFW, BGB, VKH, AMP, LMB, ESB, MSW, IJL, VVP

## Data Availability

All data, code, and materials used in the analysis are available at https://doi.org/10.6084/m9.figshare.25343779.

## Supplemental Materials

### 1. Summary of previous work on Sierra Nevada mountain chickadee study system

The hypothesis presented in the current paper is based on established results in this system, all collected over multiple years and including different groupings of birds:

1. Survival of juveniles in their first year of life is dependent on their spatial cognitive abilities, representing directional selection (40).
2. Life span of chickadees depends on their spatial cognitive abilities - birds with better spatial cognitive abilities live longer (41).
3. Spatial cognitive performance does not change with age (40, 57).
4. Spatial cognitive abilities do not show age-related senescence (49).
5. Spatial cognitive abilities are not associated with social dominance status (46) or with naturally occurring developmental perturbations (91).
6. Individual variation in spatial cognitive abilities is highly heritable and is associated with differences in genes related to cognition and brain development (47, 48).
7. Genotypes of birds with the worst cognition disappear from populations before winter (e.g., homozygous recessive birds), likely due to mortality, yet these genotypes must have been present at fledging (47, 48), which shows that birds with worse cognition die early on in their first winter.
8. There is no detectable senescence in reproduction within the naturally occurring lifespan of chickadees and females increase their reproductive success with age (73).
9. Female cognition is not associated with the number of fledglings (42) or fledgling mass (current study), however, in a high resource year, male cognition is associated with larger clutch and brood size in their social nest - a finding driven by female behavior (42).
10. Chickadees are socially monogamous species and form long-term social pairs - divorces are rare (92). In addition, it is well established that females leave their territories and visit neighboring male territories for extra-pair copulations, which we confirm here, and forced copulation does not occur in chickadees.
11. Social pairing is not associated with spatial learning and memory (43), likely because chickadees form social pairs in Fall and new recruits do not have the choice of any mate - many males are already paired (43, 92), as such the primary way to increase indirect fitness is via extra-pair copulation.
12. There are no differences in spatial cognitive abilities between males and females (80), as both rely on spatial cognition to survive the winter.
13. We have shown that supplementary food does not affect reproductive success in our system (93).

### 2. SUPPLEMENTAL RESULTS

Across 3 years of data collection on extra-pair copulations and reproductive output, we see that on average, there are 1.7 extra-pair young per nest, however, if we constrain this analysis to only nests with at least one extra-pair young, the mean is 2.5 extra-pair young per nest. Nests ranged from 0% extra-pair young to 100% extra-pair young, with the most being 6/6 nestlings sired by extra-pair fathers.

#### Elevation by year comparisons

Elevation by year comparison of extra-pair rates using binomial generalized mixed effects modeling revealed a significant interaction between elevation and year, suggesting the direction of the differences in extra-pair paternity rate between the high and low elevation sites differed across years (Figure S1; see Table S1 for sample sizes by year and elevation). Therefore, we ran a separate model for each year. In 2018, the extra-pair paternity rate was higher at the low elevation site, but this difference was not significant (elevation term: χ^2^_1_=2.24, P=0.134, N=209 offspring (134 at high, 75 at low)). In 2020, the extra-pair paternity rate was higher at the high elevation site, but this difference was not statistically significant (elevation term: χ^2^_1_=2.45, P=0.118, N=263 offspring (119 at high, 144 at low)). In 2021, the extra-pair paternity rate was higher at the low elevation site, but this difference was not statistically significant (elevation term: χ^2^_1_=1.72, P=0.190, N=260 offspring (141 at high, 119 at low)).

#### Does male age affect extra-pair paternity?

Male age was not associated with the number of extra-pair young a male sired, both according to a zero-inflated model including males that sired no extra-pair young (Age as fixed effect in the first 20 trials model: χ^2^_1_=0.83, P=0.362; Age as fixed effect in the entire task model: χ^2^_1_=2.41, P=0.121, N=280 (137 individual males)), and using a model only including males that sired at least one extra-pair young (Age as fixed effect in the first 20 trials model: χ^2^_1_=1.29, P=0.255; Age as fixed effect in the entire task model: χ^2^_1_=0.16, P=0.687, N=41nests; 34 individual males).

Male age did not affect the likelihood of a male being cuckolded (Offspring level: Age as fixed effect in the first 20 trials model: χ^2^_1_=0.08, P=0.783, Age as fixed effect in the entire task model: χ^2^_1_=0.18, P=0.676, N=348 offspring for 42 males; Nest level: Age as fixed effect in the first 20 trials model: χ^2^_1_=0.09, P=0.768; Age as fixed effect in the entire task model: χ^2^_1_=2.02, P=0.570, N=59 nests for 42 males). Within nests that had at least one extra-pair young, social male age did not predict the number of extra-pair young in the nest (χ^2^_1_=0.23, P=0.635, N=41 nests; 35 individual males).

### 3. SUPPLEMENTAL FIGURES

**Figure S1.**
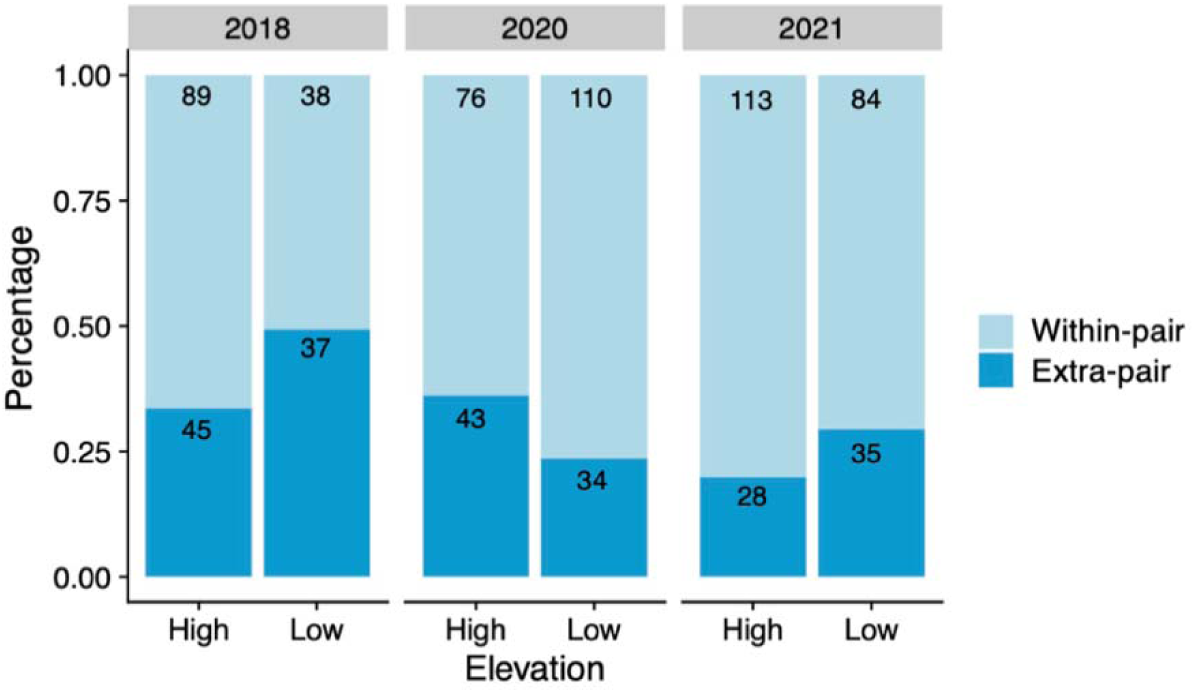
Extra-pair (EPY) and within-pair (WPY) paternity rates across 3 years of data collection and two elevation sites. Number insets represent the number of EPY and WPY for each year and elevation. EPY rates did not differ significantly by year or elevation.

**Figure S2.**
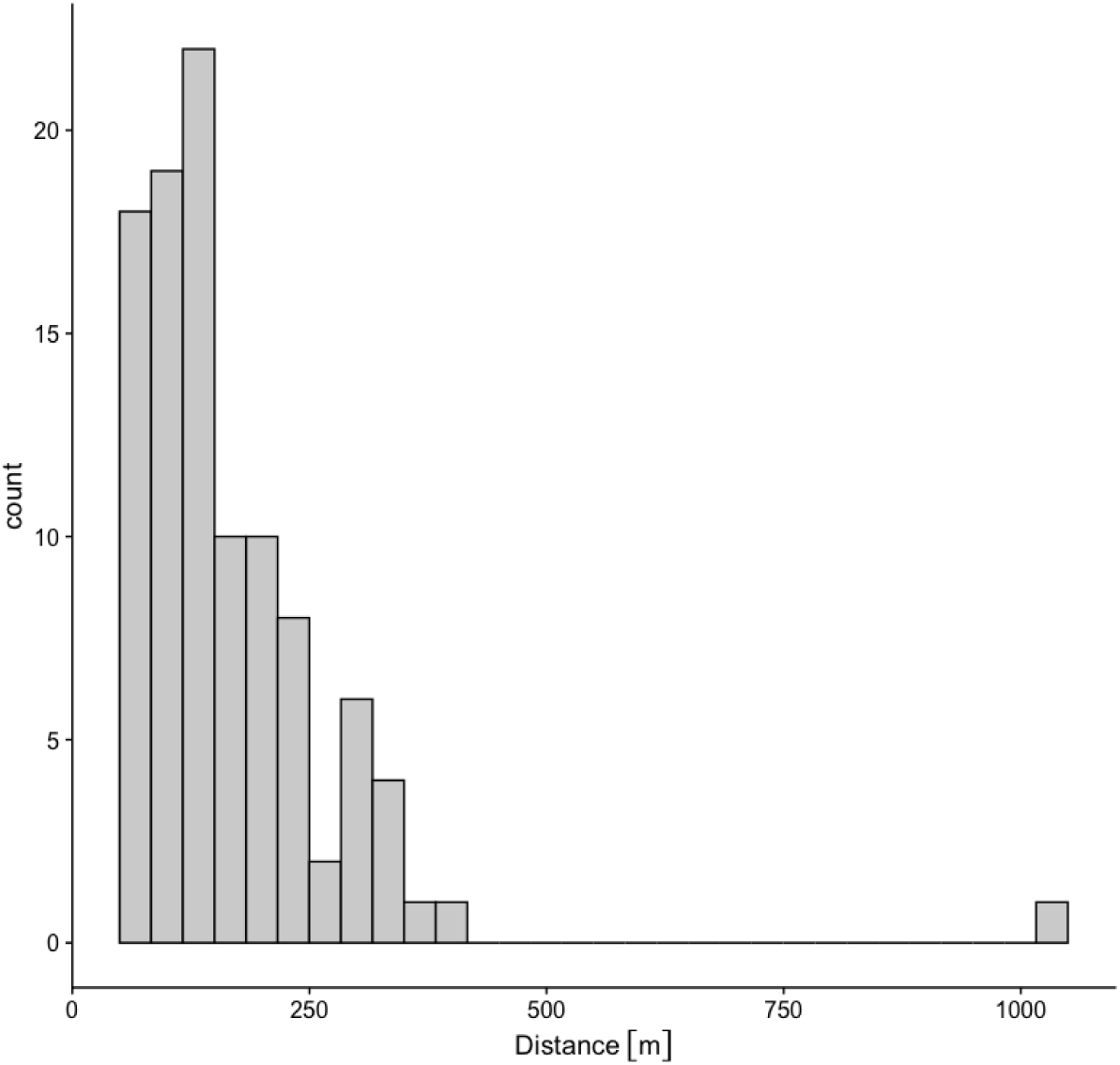
Histogram plotting the distance of extra-pair male nests from the females’ nest. Mean = 167 meters and Median = 136 meters. Range is 56 to 1022 meters between nestboxes.

**Figure S3a.**
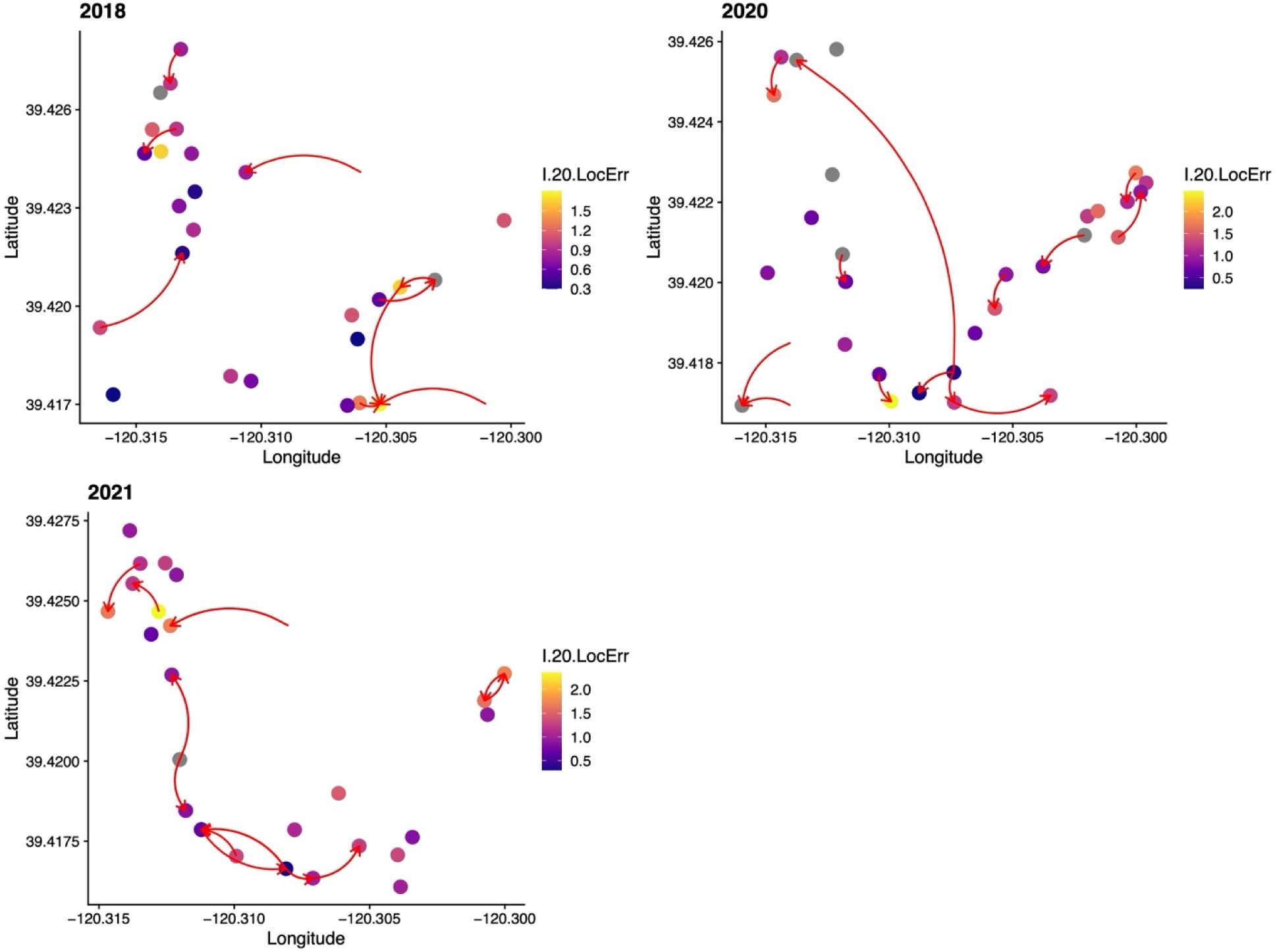
High elevation extra-pair paternity relationships and cognition scores. Maps show nests with genotyped offspring and the nests of males that sired extra-pair offspring. Red arrows show extra-pair paternity relationships. The direction of the arrow indicates a social male’s nest at the blunt end connected to the nest he sired extra-pair young in at the sharp end. Arrows that originate in open space show males that sired extra-pair young but did not breed in our nest boxes in that year (they likely bred in a natural cavity). Point colors represent the cognition scores of the social male at each nest. Gray circles represent males without cognition data. Only nests that included EPY are included on this map, which is a sample of the active nests in each year (Figure S4).

**Figure S3b.**
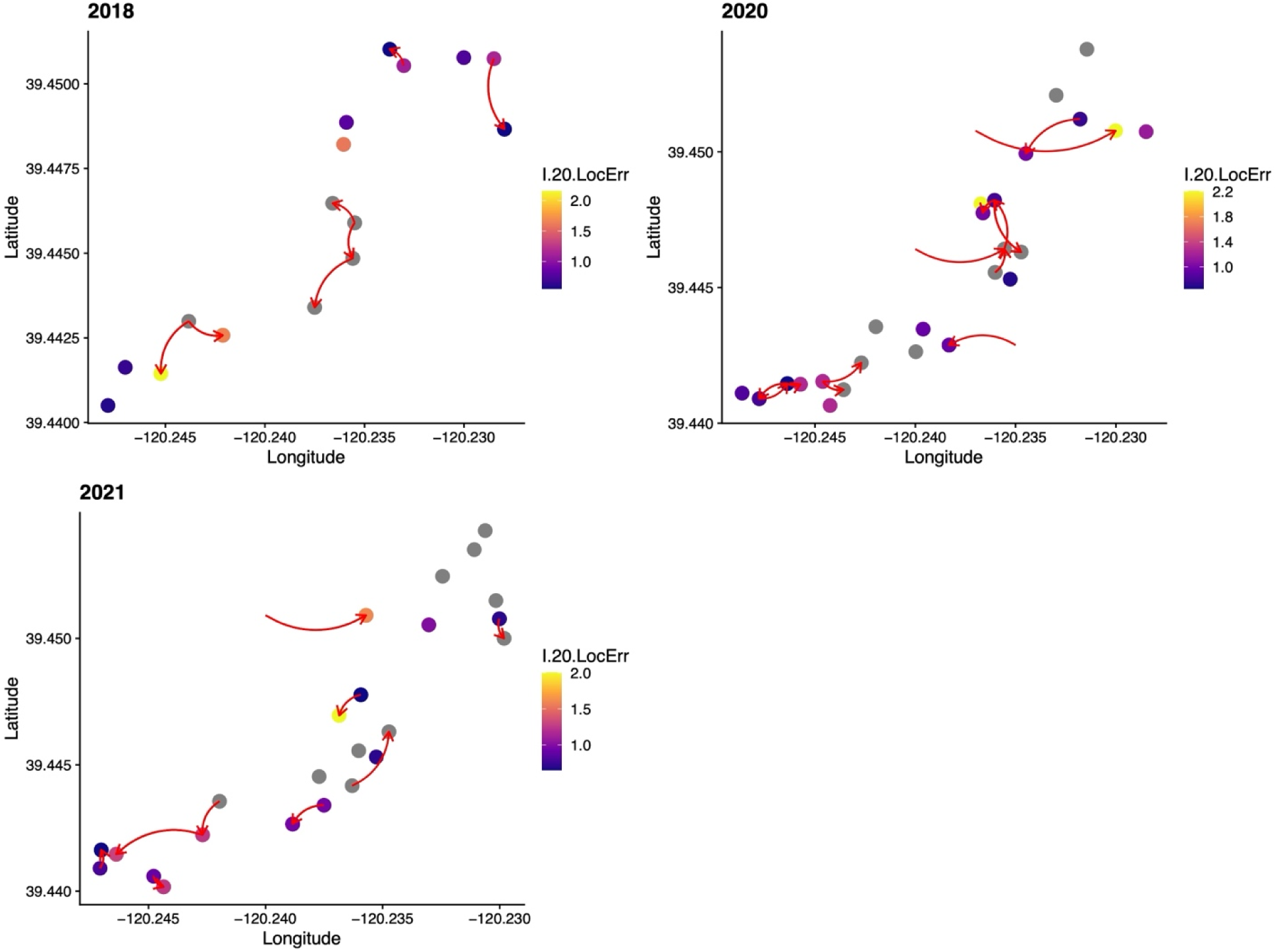
Low elevation extra-pair paternity relationships and cognition scores. Maps show nests with genotyped offspring and the nests of males that sired extra-pair offspring. Red arrows show extra-pair paternity relationships. The direction of the arrow indicates a social male’s nest at the blunt end connected to the nest he sired extra-pair young in at the sharp end. Arrows that originate in open space show males that sired extra-pair young but did not breed in our nest boxes in that year (they likely bred in a natural cavity). Point colors represent the cognition scores of the social male at each nest. Gray circles represent males without cognition data. Only nests that included EPY are included on this map, which is a sample of the active nests in each year (Figure S4).

**Figure S4.**
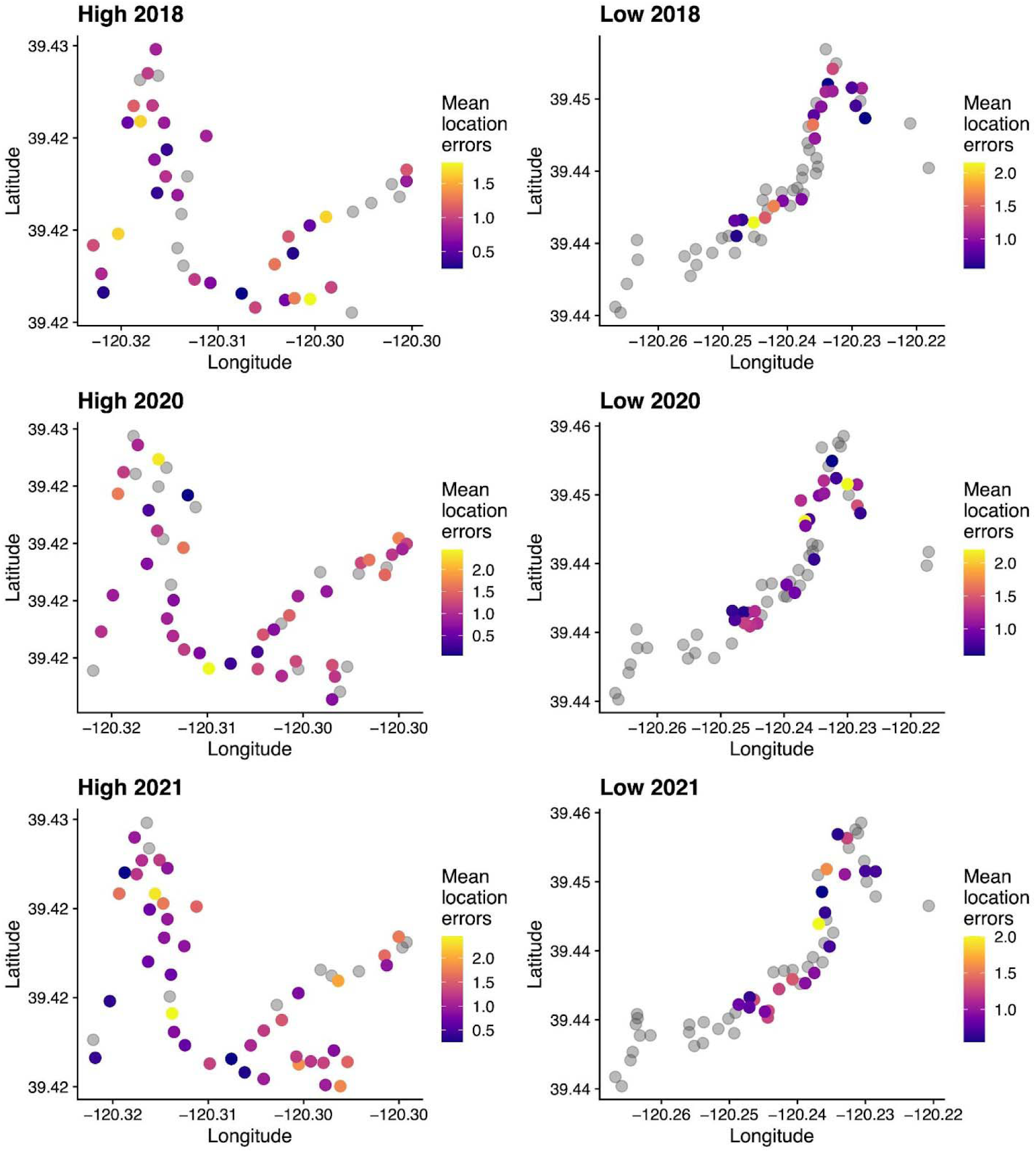
Heat maps of male spatial cognitive scores for high and low elevation nestboxes across the 3 years we have extra-pair paternity data from, including those nests that failed before day 16 or were not genotyped. Each point represents a nestbox. Color gradient represents mean location errors of male on the first 20 trials of the spatial cognitive task. Gray circles indicate nestboxes with no cognitive data for the male. Data shows that nestboxes are not clustered by male cognitive performance. Some of the males on the maps did not participate in cognitive testing until after these breeding seasons, but since spatial cognitive ability does not change with age, we present their scores here as well.

**Figure S5.**
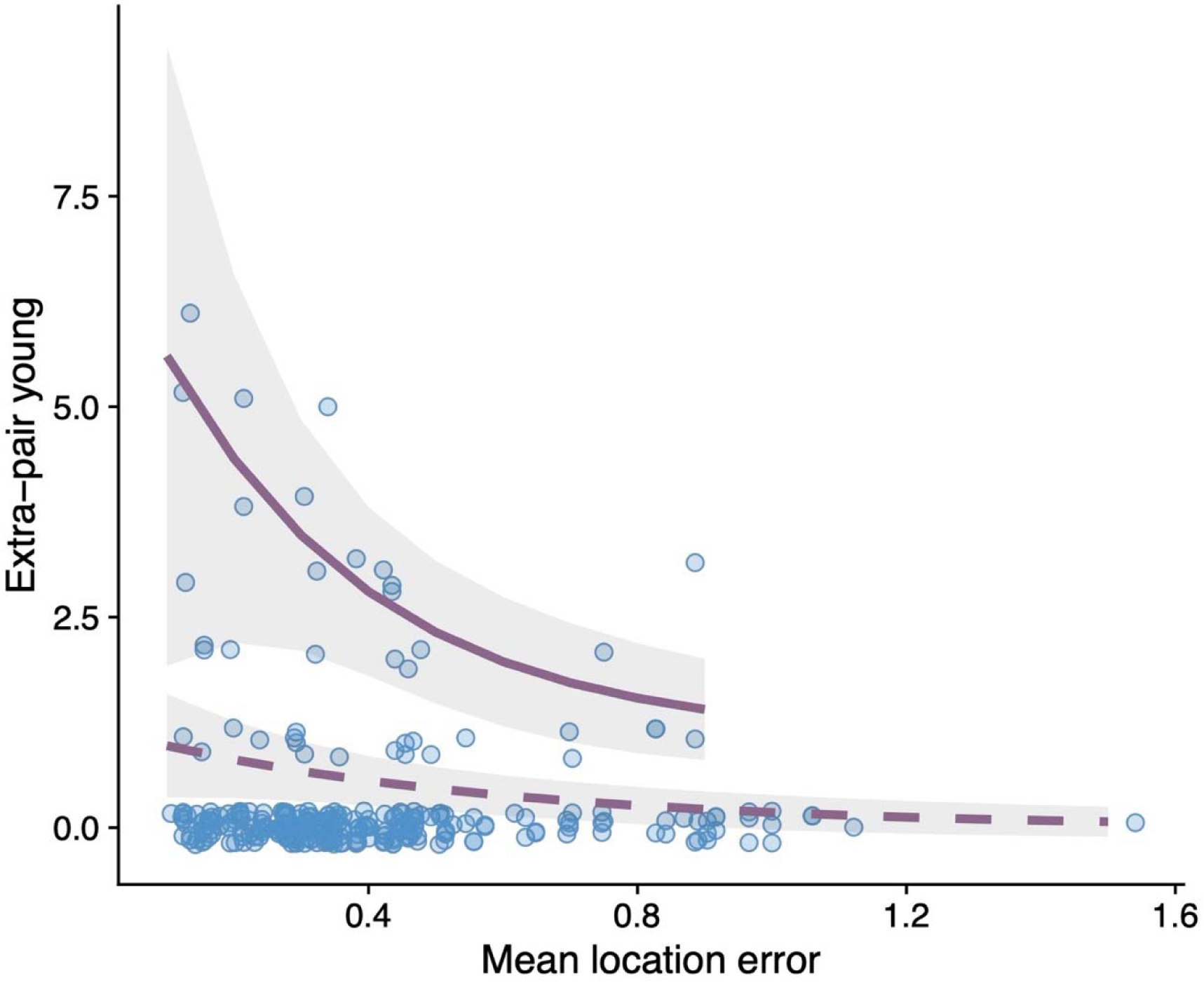
Model predictions of the number of extra-pair young a male sired in a year by the mean number of location errors per trial over the 4-day testing period. The dashed line represents a zero-inflated model for all males, including those with no extra-pair young (N=280; 137 individual males), and the solid line represents the model prediction for males that sired at least one extra-pair young (N=41; 34 individual males). Shading represents bootstrapped 95% confidence intervals. Note: data presented in figure do not account for total trials completed, which was accounted for in statistical analysis.

**Figure S6.**
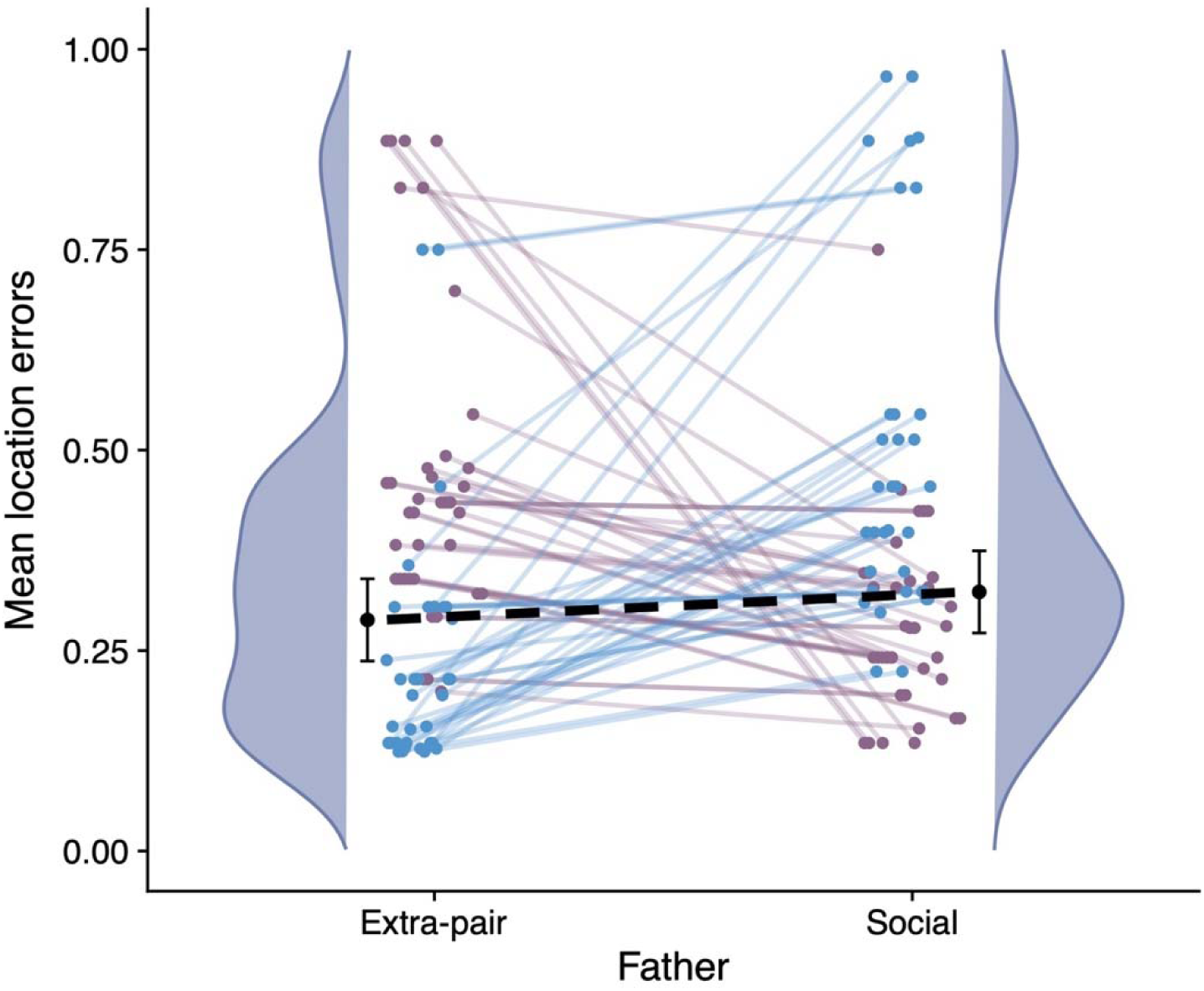
Comparing spatial cognitive performance (mean number of location errors per trial over the 4-day task) of extra-pair and social fathers (total of 71 comparisons). Lines connect males by offspring: blue lines show comparisons where the extra-pair father performed better than the social father (34 of 71 cases or 47.9%), and purple lines show comparisons where the social father performed better than the extra-pair father (37 of 71 cases or 52.1%). Shaded distributions represent smoothed histograms of cognitive performance. Note that the data presented in the figure do not account for the total number of trials completed, which was accounted for in the statistical analysis.

**Figure S7a.**
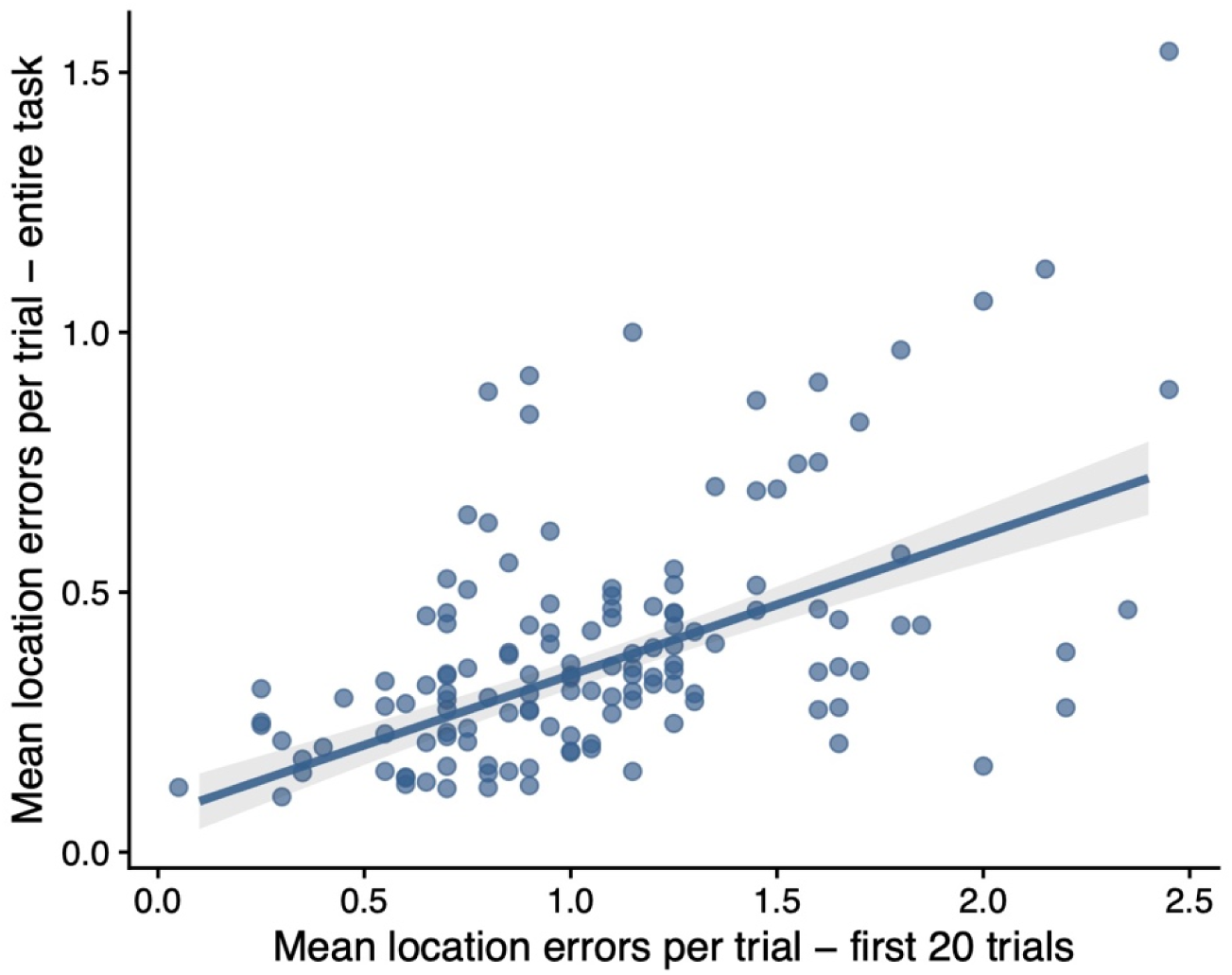
There is a strong positive correlation between the number of location errors on the entire 4-day task and on the first 20 trials for males completing the task (Coef=0.27, F=121.72, P<0.001).

**Figure S7b.**
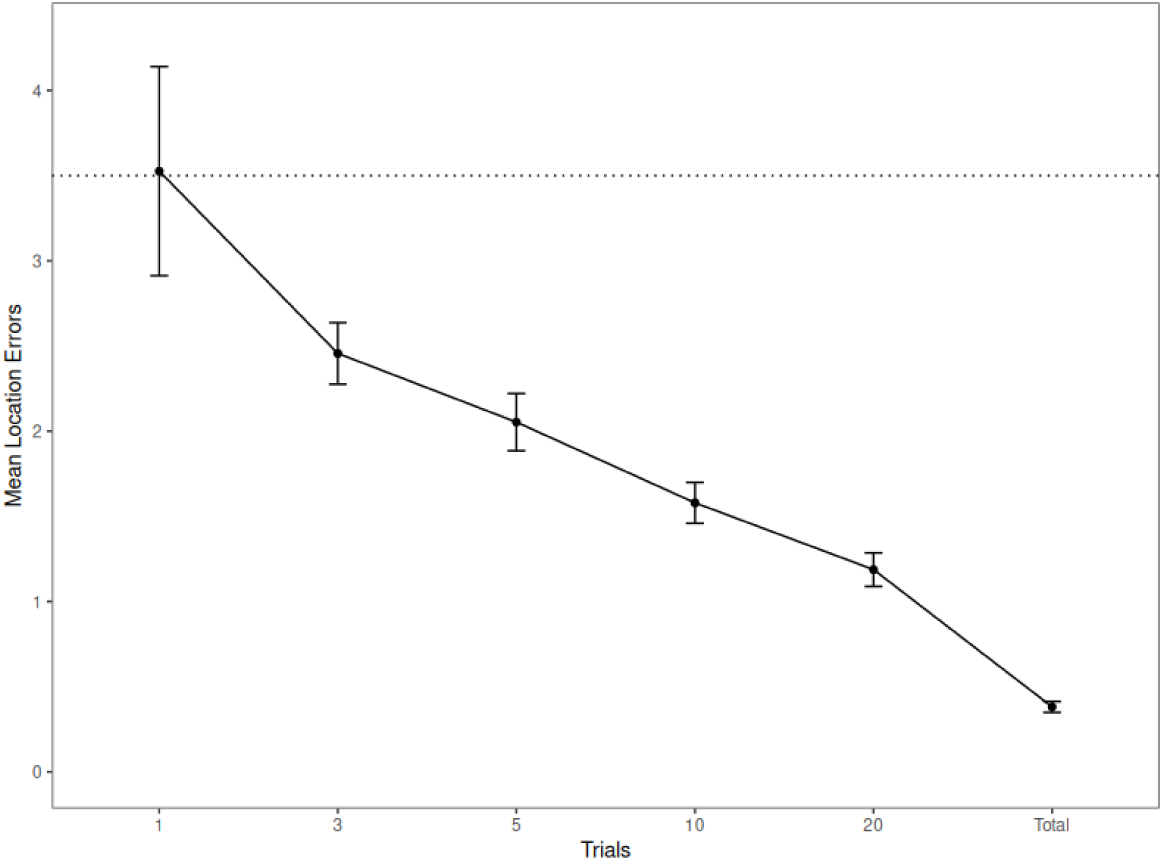
Representative learning curve of 19 males included in the extra-pair paternity analysis, from the 2021 breeding season. Dotted line represented chance level performance on the task at 3.5 mean location errors. Curve includes performance on trials 1 – 20 and over the entire 4-day task. Errors bars represent standard error of the mean.

**Figure S8.**
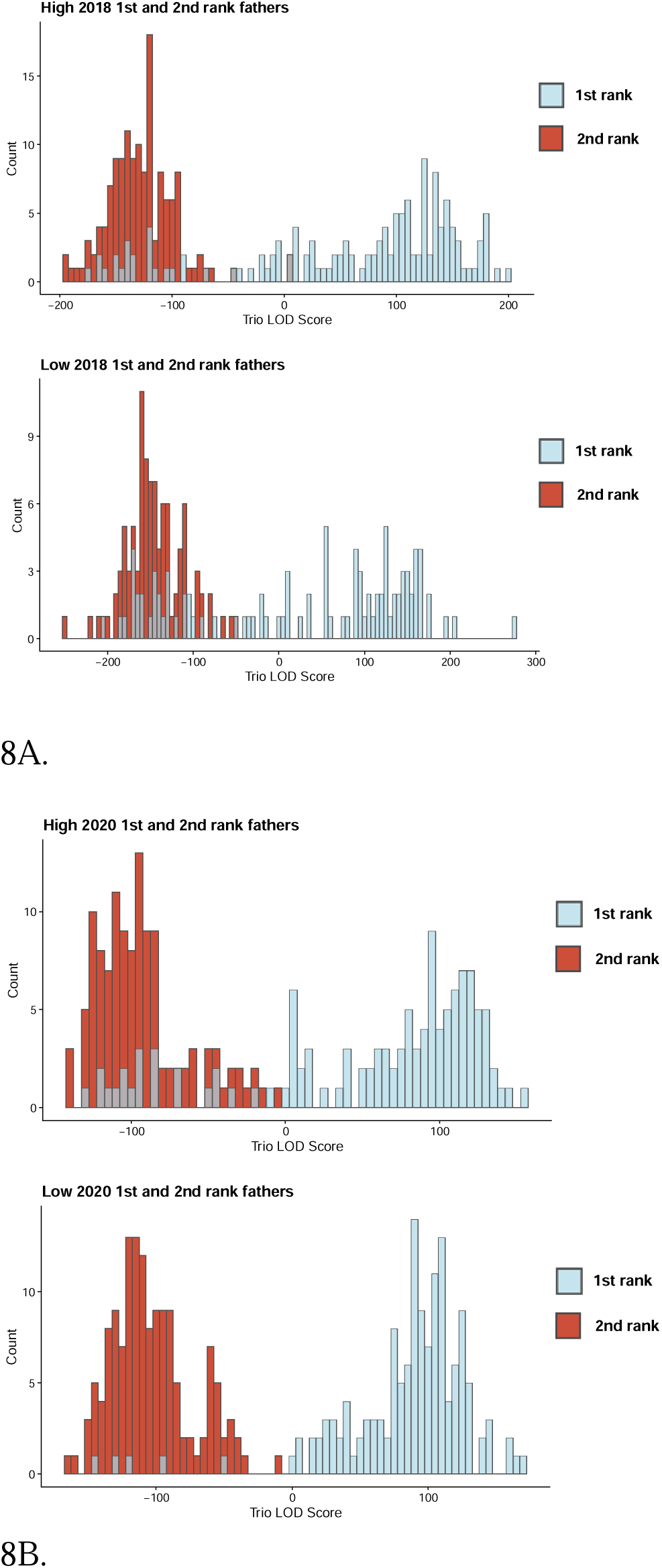
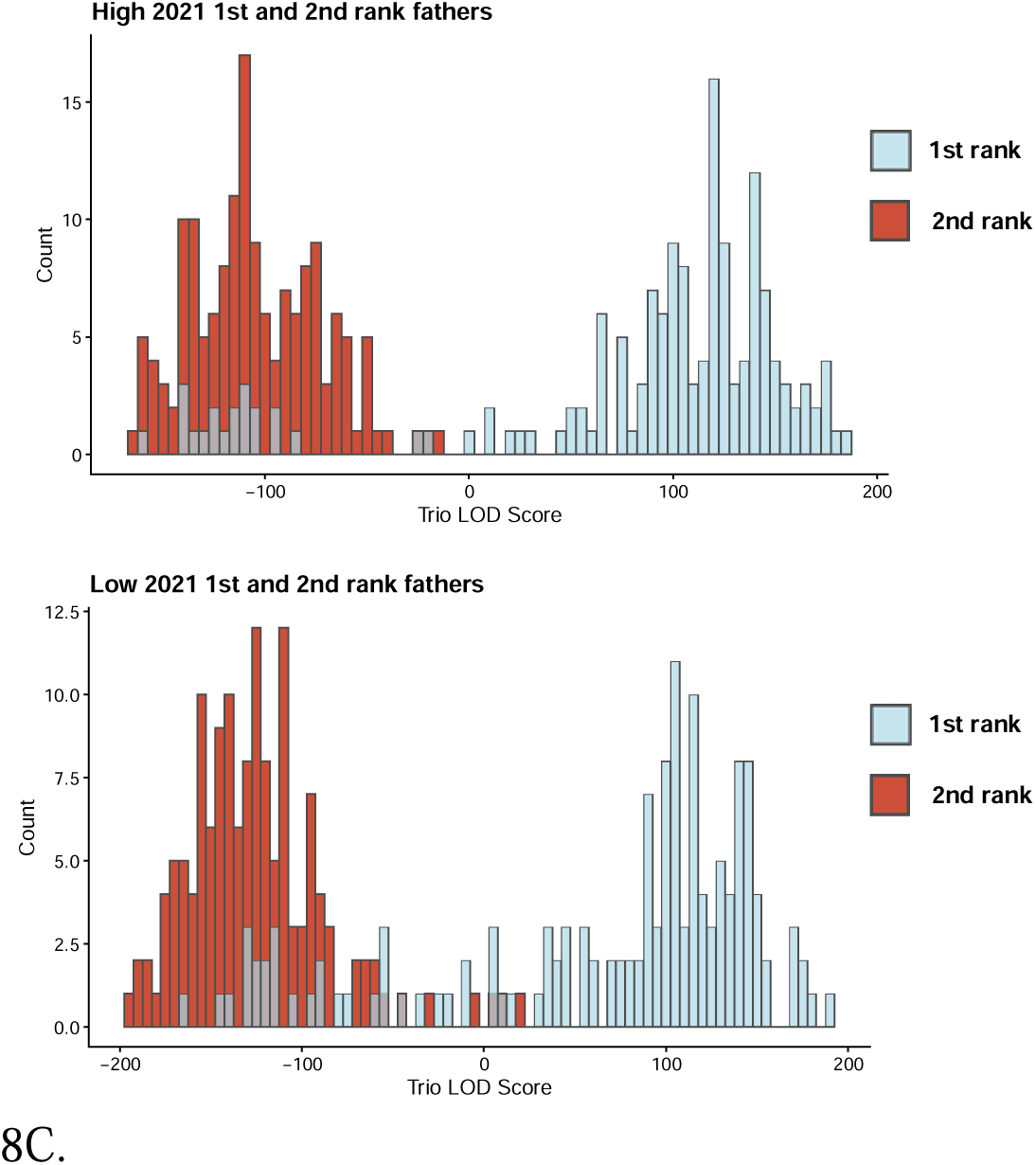
Comparisons of Trio LOD scores for 1st and 2nd ranked potential fathers for each offspring by elevation and year. Note the mostly bimodal distribution in which 1st-ranked males (blue) consistently out-performed 2nd-ranked males (red). Offspring with low 1st-ranked male Trio LOD scores are cases where we did not sample the offspring’s genetic father.

### 4. SUPPLEMENTAL TABLES

**Table S1.** Sample sizes for extra-pair paternity broken down by year and elevation for both the nest and offspring level.

|  | Year | Elevation | No EPY | ≥1 EPY | Rate |
| --- | --- | --- | --- | --- | --- |
| Nest level |  |  |  |  |  |
|  | 2018 | High | 8 | 17 | 68.00% |
|  |  | Low | 1 | 13 | 92.90% |
|  | 2020 | High | 6 | 18 | 75.00% |
|  |  | Low | 8 | 14 | 63.60% |
|  | 2021 | High | 9 | 13 | 59.10% |
|  |  | Low | 6 | 14 | 70.00% |
|  | Total | High | 23 | 48 | 67.60% |
|  |  | Low | 15 | 41 | 73.20% |
| Offspring level |  |  |  |  |  |
|  | Year | Elevation | WPY | EPY | Rate |
|  | 2018 | High | 89 | 45 | 33.60% |
|  |  | Low | 38 | 37 | 49.30% |
|  | 2020 | High | 76 | 43 | 36.10% |
|  |  | Low | 110 | 34 | 23.60% |
|  | 2021 | High | 113 | 28 | 19.90% |
|  |  | Low | 84 | 35 | 29.40% |
|  | Total | High | 278 | 116 | 29.40% |
|  |  | Low | 232 | 106 | 31.40% |

**Table S2.** Differences in extra-pair paternity rates between high and low elevations. A likelihood ratio test indicated that an interaction between elevation and year improved the model, so years were run separately.

|  | Estimate | Standard error | Z value |
| --- | --- | --- | --- |
| Intercept | -0.85 | 0.35 | -2.42 |
| Elevation (Low) | 0.84 | 0.56 | 1.50 |
| Random effects | Variance | Std deviation |  |
| Mother ID | 1.66 | 1.29 |  |

N=209 offspring, 39 mothers
|  | Estimate | Standard error | Z value |
| --- | --- | --- | --- |
| Intercept | -0.72 | 0.39 | -1.86 |
| Elevation (Low) | -0.88 | 0.56 | -1.57 |
| Random effects | Variance | Std deviation |  |
| Mother ID | 2.15 | 1.47 |  |

N=263 offspring, 46 mothers
|  | Estimate | Standard error | Z value |
| --- | --- | --- | --- |
| Intercept | -1.79 | 0.40 | -4.53 |
| Elevation (Low) | 0.69 | 0.53 | 1.31 |
| Random effects | Variance | Std deviation |  |
| Mother ID | 1.56 | 1.25 |  |

N=260 offspring, 42 mothers

**Table S3.** Relationship between spatial cognitive ability and extra-pair young sired. Two models are presented for each spatial cognitive measure, a model that included a zero-inflation parameter for all observations and a model that only included male-years where the male sired at least one offspring.

**3A. First 20 trials mean location error with zero-inflation parameter**
| Conditional model | Estimate | Standard error | Z value |
| --- | --- | --- | --- |
| Intercept | 0.40 | 0.54 | 0.75 |
| First 20 trials mean location error | -0.92 | 0.37 | -2.51 |
| Elevation (Low) | 0.61 | 0.31 | 1.99 |
| Year 2020 | 0.67 | 0.43 | 1.54 |
| Year 2021 | 0.74 | 0.43 | 1.70 |
| Zero-inflation model | Estimate | Standard error | Z value |
| Intercept | 1.28 | 0.23 | 5.55 |
| Random effects | Variance | Std deviation |  |
| Male ID | 0.08 | 0.29 |  |

**3B. First 20 trials mean location error with positive-only values**
|  | Estimate | Standard error | Z value |
| --- | --- | --- | --- |
| Intercept | 0.50 | 0.63 | 0.79 |
| First 20 trials mean location error | -1.16 | 0.48 | -2.44 |
| Elevation (Low) | 0.74 | 0.34 | 2.17 |
| Year 2020 | 0.73 | 0.55 | 1.33 |
| Year 2021 | 0.76 | 0.55 | 1.38 |
| Random effects | Variance | Std deviation |  |
| Male ID | 0.11 | 0.33 |  |

**3C. Entire task mean location error with zero-inflation parameter**
| Conditional model | Estimate | Standard error | Z value |
| --- | --- | --- | --- |
| Intercept | 0.18 | 0.46 | 0.40 |
| Entire task mean location error | -1.86 | 0.65 | -2.85 |
| Elevation (Low) | 0.76 | 0.29 | 2.65 |
| Year 2020 | 0.73 | 0.42 | 1.75 |
| Year 2021 | 0.79 | 0.41 | 1.91 |
| Zero-inflation model | Estimate | Standard error | Z value |
| Intercept | 1.34 | 0.22 | 6.07 |
| Random effects | Variance | Std deviation |  |
| Male ID | 0.03 | 0.17 |  |

N=280 male-years (137 unique males over 3 breeding seasons)
| 3D. Entire task mean location error with positive-only values |  |  |  |
| --- | --- | --- | --- |
|  | Estimate | Standard error | Z value |
| Intercept | 0.24 | 0.56 | 0.43 |
| Entire task mean location error | -2.55 | 0.92 | -2.77 |
| Elevation (Low) | 0.85 | 0.32 | 2.65 |
| Year 2020 | 0.81 | 0.54 | 1.50 |
| Year 2021 | 0.88 | 0.54 | 1.64 |
| Random effects | Variance | Std deviation |  |
| Male ID | 0.03 | 0.18 |  |

N=41 male-years (34 unique males over 3 breeding seasons)

**Table S4.** Relationship between male and female spatial cognitive ability and breeding performance. Results calculated using mean location error over the entire task as fixed effects may be less informative than models using the first 20 trials scores as fixed effects. The total number of trials completed over the entire task is a strong predictor of mean location errors over the entire task, but when mean location errors over the entire task is included as a fixed effect (on the right side of the equation), we could not control for the number of total trials completed over the entire task.

4A. Brood size: first 20 trials
|  | Estimate | Standard error | Z value |
| --- | --- | --- | --- |
| Intercept | 1.75 | 0.07 | 26.07 |
| Male First 20 trials mean location error | 0.06 | 0.04 | 1.52 |
| Female First 20 trials mean location error | -0.05 | 0.04 | -1.32 |
| Elevation (Low) | 0.08 | 0.04 | 2.10 |
| Random effects | Variance | Std deviation |  |
| Year (2015-2023) | <0.01 | 0.08 |  |
| Male ID | <0.01 | 0.09 |  |
| Female ID | <0.01 | 0.06 |  |

4B. Brood size: entire task
|  | Estimate | Standard error | Z value |
| --- | --- | --- | --- |
| Intercept | 1.75 | 0.06 | 29.78 |
| Male mean location error over entire task | 0.04 | 0.10 | 0.40 |
| Female mean location error over entire task | -0.04 | 0.09 | -0.48 |
| Elevation (Low) | 0.09 | 0.04 | 2.32 |
| Random effects | Variance | Std deviation |  |
| Year (2015-2023) | 0.01 | 0.07 |  |
| Male ID | 0.01 | 0.10 |  |
| Female ID | <0.01 | 0.05 |  |

4C. Brood size relative to clutch size: first 20 trials
|  | Estimate | Standard error | Z value |
| --- | --- | --- | --- |
| Intercept | 0.72 | 0.10 | 7.39 |
| Male First 20 trials mean location error | 0.02 | 0.02 | 1.00 |
| Female First 20 trials mean location error | 0.02 | 0.03 | 0.70 |
| Elevation (Low) | 0.02 | 0.02 | 0.97 |
| Clutch size | 0.14 | 0.01 | 12.05 |
| Random effects | Variance | Std deviation |  |
| Year (2015-2023) | <0.01 | 0.04 |  |
| Male ID | <0.01 | <0.01 |  |
| Female ID | <0.01 | <0.01 |  |

**4D. Brood size relative to clutch size: entire task**
|  | Estimate | Standard error | Z value |
| --- | --- | --- | --- |
| Intercept | 0.74 | 0.09 | 7.93 |
| Male mean location error over entire task | <0.01 | 0.06 | 0.03 |
| Female mean location error over entire task | 0.05 | 0.06 | 0.88 |
| Elevation (Low) | 0.02 | 0.02 | 1.03 |
| Clutch size | 0.15 | 0.01 | 12.36 |
| Random effects | Variance | Std deviation |  |
| Year (2015-2023) | <0.01 | 0.04 |  |
| Male ID | <0.01 | <0.01 |  |
| Female ID | <0.01 | <0.01 |  |

**4E. Mean nestling mass: first 20 trials**
|  | Estimate | Standard error | Df | T value |
| --- | --- | --- | --- | --- |
| Intercept | 12.47 | 0.25 | 67.48 | 50.35 |
| Male first 20 trials mean location error | -0.28 | 0.13 | 94.91 | -2.20 |
| Female first 20 trials mean location error | 0.15 | 0.15 | 46.89 | 0.97 |
| Elevation (Low) | -0.38 | 0.13 | 56.62 | -2.86 |
| Random effects | Variance | Std deviation |  |  |
| Year (2015-2023) | 0.10 | 0.31 |  |  |
| Male ID | 0.04 | 0.19 |  |  |
| Female ID | 0.13 | 0.35 |  |  |
| Residual | 0.41 | 0.64 |  |  |

**4F. Mean nestling mass: entire task**
|  | Estimate | Standard error | Df | T value |
| --- | --- | --- | --- | --- |
| Intercept | 12.37 | 0.22 | 54.65 | 56.37 |
| Male mean location error over entire task | -0.19 | 0.33 | 131.24 | -0.57 |
| Female mean location error over entire task | 0.11 | 0.32 | 70.54 | 0.33 |
| Elevation (Low) | -0.40 | 0.13 | 54.32 | -3.00 |
| Random effects | Variance | Std deviation |  |  |
| Year (2015-2023) | 0.10 | 0.31 |  |  |
| Male ID | 0.06 | 0.25 |  |  |
| Female ID | 0.10 | 0.32 |  |  |
| Residual | 0.41 | 0.64 |  |  |

N=202 nests over 9 years for 107 males and 89 females.

**Table S5.** Test of whether siring extra-pair offspring affected the offspring in a male’s own nest. Whether a male sired extra-pair offspring or not was compared to A) brood size, B) brood size controlled for clutch size, and C) mean nestling mass.

5A. Brood size
|  | Estimate | Standard error | Z value |
| --- | --- | --- | --- |
| Intercept | 1.78 | 0.04 | 45.66 |
| Sired EPY (Yes) | 0.03 | 0.05 | 0.55 |
| Elevation (Low) | 0.04 | 0.04 | 0.95 |
| Year 2020 | -0.06 | 0.04 | -1.33 |
| Year 2021 | 0.06 | 0.04 | 1.34 |
| Random effects | Variance | Std deviation |  |
| Male ID | 0.01 | 0.12 |  |
| Female ID | <0.01 | 0.04 |  |

5B. Brood size relative to clutch size
|  | Estimate | Standard error | Z value |
| --- | --- | --- | --- |
| Intercept | 0.71 | 0.10 | 7.09 |
| Sired EPY (Yes) | <0.01 | 0.03 | 0.03 |
| Elevation (Low) | <-0.01 | 0.03 | -0.17 |
| Clutch size | 0.15 | 0.01 | 11.41 |
| Year 2020 | -0.03 | 0.03 | -0.86 |
| Year 2021 | -0.01 | 0.03 | -0.26 |
| Random effects | Variance | Std deviation |  |
| Male ID | <0.01 | <0.01 |  |
| Female ID | <0.01 | <0.01 |  |

5C. Mean nestling mass
|  | Estimate | Standard error | Df | T value |
| --- | --- | --- | --- | --- |
| Intercept | 12.41 | 0.14 | 134.89 | 87.98 |
| Sired EPY (Yes) | 0.26 | 0.18 | 134.89 | 1.45 |
| Elevation (Low) | -0.32 | 0.16 | 85.75 | -1.94 |
| Year 2020 | -0.68 | 0.17 | 115.82 | -3.94 |
| Year 2021 | -0.14 | 0.17 | 115.13 | -0.82 |
| Random effects | Variance | Std deviation |  |  |
| Male ID | 0.12 | 0.35 |  |  |
| Female ID | 0.00 | 0.00 |  |  |
| Residual | 0.61 | 0.78 |  |  |

N=140 nests for 98 males and 101 females

**Table S6.**
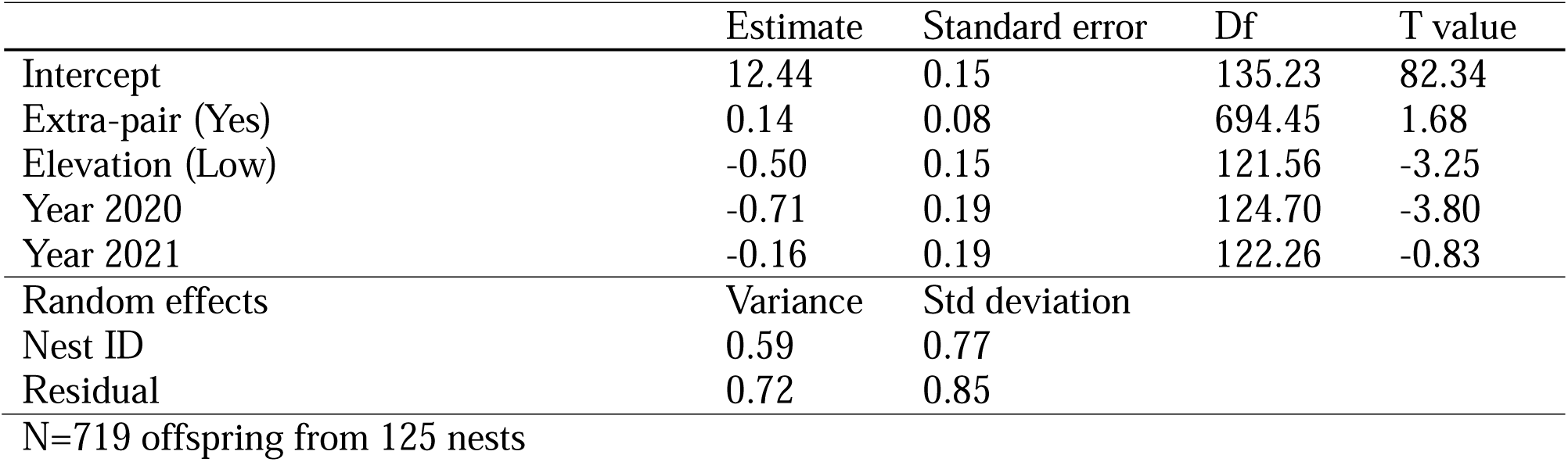
Test of whether offspring sired by extra-pair fathers differed in mass from offspring sired by social males.

|  | Estimate | Standard error | Df | T value |
| --- | --- | --- | --- | --- |
| Intercept | 12.44 | 0.15 | 135.23 | 82.34 |
| Extra-pair (Yes) | 0.14 | 0.08 | 694.45 | 1.68 |
| Elevation (Low) | -0.50 | 0.15 | 121.56 | -3.25 |
| Year 2020 | -0.71 | 0.19 | 124.70 | -3.80 |
| Year 2021 | -0.16 | 0.19 | 122.26 | -0.83 |
| Random effects | Variance | Std deviation |  |  |
| Nest ID | 0.59 | 0.77 |  |  |
| Residual | 0.72 | 0.85 |  |  |

N=719 offspring from 125 nests

**Table S7.** Comparison of spatial cognition scores between social males and extra-pair males they were cuckolded by. Log-transformed values are reported in Table 4B – entire task mean location error model.

**7A. First 20 trials mean location error model**
|  | Estimate | Standard error | Df | T value |
| --- | --- | --- | --- | --- |
| Intercept | 1.00 | 0.09 | 37.00 | 11.21 |
| Father type (social) | 0.21 | 0.05 | 108.11 | 3.85 |
| Elevation (Low) | -0.10 | 0.13 | 27.36 | -0.73 |
| Random effects | Variance | Std deviation |  |  |
| Offspring ID | 0.00 | 0.00 |  |  |
| Mother ID | 0.10 | 0.32 |  |  |
| Residual | 0.11 | 0.32 |  |  |

**7B. Entire task mean location error model**
|  | Estimate | Standard error | Df | T value |
| --- | --- | --- | --- | --- |
| Intercept | -1.07 | 0.06 | 42.35 | -16.78 |
| Father type (social) | 0.11 | 0.05 | 107.68 | 2.44 |
| Elevation (Low) | -0.17 | 0.09 | 27.30 | -1.93 |
| Trials completed | -4.80 | 0.33 | 136.81 | -14.63 |
| Trials completed<sup>2</sup> | 1.67 | 0.33 | 136.51 | 5.03 |
| Random effects | Variance | Std deviation |  |  |
| Offspring ID | 0.00 | 0.00 |  |  |
| Mother ID | 0.04 | 0.19 |  |  |
| Residual | 0.08 | 0.28 |  |  |

N=71 social male – extra-pair male comparisons (71 offspring) from 32 mothers

**Table S8.** Test of whether a male’s spatial cognition score was associated with whether he was cuckolded or not.

**8A. First 20 trials mean location error offspring model**
|  | Estimate | Standard error | Z value |
| --- | --- | --- | --- |
| Intercept | -2.05 | 0.75 | -2.74 |
| Male first 20 trials mean location error | -0.20 | 0.40 | -0.50 |
| Female first 20 trials mean location error | 1.23 | 0.49 | 2.49 |
| Elevation (Low) | 0.52 | 0.41 | 1.26 |
| Year 2020 | -0.83 | 0.38 | -2.15 |
| Year 2021 | -0.55 | 0.39 | -1.40 |
| Random effects | Variance | Std deviation |  |
| Male ID | 0.23 | 0.48 |  |
| Female ID | 0.41 | 0.64 |  |

**8B. Entire task mean location error offspring model**
|  | Estimate | Standard error | Z value |
| --- | --- | --- | --- |
| Intercept | -0.79 | 0.61 | -1.31 |
| Male entire task mean location error | -0.67 | 1.02 | -0.65 |
| Female entire task mean location error | 0.41 | 0.97 | 0.42 |
| Elevation (Low) | 0.16 | 0.44 | 0.37 |
| Year 2020 | -0.81 | 0.40 | -2.02 |
| Year 2021 | -0.53 | 0.40 | -1.32 |
| Random effects | Variance | Std deviation |  |
| Male ID | 0.05 | 0.22 |  |
| Female ID | 0.86 | 0.93 |  |

**8C. First 20 trials mean location error nest model**
|  | Estimate | Standard error | Z value |
| --- | --- | --- | --- |
| Intercept | 0.35 | 1.11 | 0.32 |
| Male first 20 trials mean location error | 0.16 | 0.67 | 0.24 |
| Female first 20 trials mean location error | 0.35 | 0.76 | 0.46 |
| Elevation (Low) | 0.20 | 0.64 | 0.30 |
| Year 2020 | 0.21 | 0.73 | 0.29 |
| Year 2021 | -0.73 | 0.72 | -1.01 |
| Random effects | Variance | Std deviation |  |
| Male ID | <0.01 | <0.01 |  |
| Female ID | <0.01 | <0.01 |  |

**8D. Entire task mean location error nest model**
|  | Estimate | Standard error | Z value |
| --- | --- | --- | --- |
| Intercept | 0.82 | 0.91 | 0.90 |
| Male entire task mean location error | 0.80 | 1.72 | 0.47 |
| Female entire task mean location error | -0.52 | 1.48 | -0.35 |
| Elevation (Low) | 0.07 | 0.62 | 0.11 |
| Year 2020 | 0.32 | 0.73 | 0.44 |
| Year 2021 | -0.71 | 0.73 | -0.98 |
| Random effects | Variance | Std deviation |  |
| Male ID | <0.01 | <0.01 |  |
| Female ID | <0.01 | <0.01 |  |

**Table S9.** Test of whether the number of extra-pair young in a nest is associated with spatial cognitive ability. Results calculated using mean location error over the entire task as fixed effects may be less informative than models using the first 20 trials scores as fixed effects. The total number of trials completed over the entire task is a strong predictor of mean location errors over the entire task, but when mean location errors over the entire task is included as a fixed effect (on the right side of the equation), we could not control for the number of total trials completed over the entire task.

9A. First 20 trials model
|  | Estimate | Standard error | Z value |
| --- | --- | --- | --- |
| Intercept | -0.26 | 0.55 | -0.48 |
| Male First 20 trials mean location error | -0.30 | 0.30 | -1.01 |
| Female First 20 trials mean location error | 0.92 | 0.35 | 2.61 |
| Elevation (Low) | 0.79 | 0.35 | 2.23 |
| Year 2020 | -0.70 | 0.34 | -2.09 |
| Year 2021 | 0.09 | 0.31 | 0.31 |
| Random effects | Variance | Std deviation |  |
| Father ID | <0.01 | <0.01 |  |
| Mother ID | <0.01 | <0.01 |  |

9B. Entire task model
|  | Estimate | Standard error | Z value |
| --- | --- | --- | --- |
| Intercept | 0.54 | 0.43 | 1.27 |
| Male mean location error over entire task | -0.42 | 0.67 | -0.63 |
| Female mean location error over entire task | 0.74 | 0.73 | 1.02 |
| Elevation (Low) | 0.33 | 0.28 | 1.19 |
| Year 2020 | -0.69 | 0.34 | -2.02 |
| Year 2021 | 0.14 | 0.30 | 0.45 |
| Random effects | Variance | Std deviation |  |
| Father ID | <0.01 | <0.01 |  |
| Mother ID | <0.01 | <0.01 |  |

## References

1. Andersson, M & Simmons, LW. Sexual selection and mate choice. Trends Ecol. Evol. 21, 296–302 (2006).

2. Rosenthal, G. *Mate Choice: the Evolution of Sexual Decision Making from Microbes to Humans*. Princeton, NJ, Princeton University Press (2017).

3. Davies, AD, Lewis, Z, & Dougherty LR. A meta-analysis of factors influencing the strength of mate-choice copying in animals. Behav. Ecol. 31, 1279–1290 (2020).

4. Henshaw, JM, Fromhage, L, & Jones, AG. The evolution of mating preferences for genetic attractiveness and quality in the presence of sensory bias. PNAS 119, e2206262119 (2022).

5. DuVal, EH, Fitzpatrick, CL, Hobson, EA, & Servedio, MR. Inferred attractiveness: A generalized mechanism for sexual selection that can maintain variation in traits and preferences over time. PLoS Biol. 21, e3002269 (2023).

6. Fisher, RA. *The genetical theory of natural selection*. Oxford, Clarendon Press (1930).

7. Prokop, Z.M., Michalczyk, £., Drobniak, S.M., Herdegen, M. and Radwan, J. Meta-analysis suggests choosy females get sexy sons more than “good genes”. Evolution. 66, 2665–2673 (2012).

8. Endler, JA. Sensory drive: does sensory biology bias or constrain the direction of evolution? Am. Nat. 139, S1–S153 (1992).

9. Andersson, M. *Sexual Selection*. Princeton, NJ, Princeton University Press (1994).

10. Bonduriansky, R. The evolution of male mate choice in insects: a synthesis of ideas and evidence. Biological Reviews. 76, 305–339 (2001).

11. Gowaty, PA, Steinichen, R, & Anderson, WW. Indiscriminate females and choosy males: within and between species variation in Drosophila. Evolution. 57, 2037–2045 (2003).

12. Verner, J, Wilson, MF. Mating systems, sexual dimorphism and the role male North American passerine birds in the nesting cycle. Oenitol. Monogr. 9, 1–76 (1969).

13. Schärer, L, Rowe, L, & Arnquist, G. Anisogamy, chance and the evolution of sex roles. Trends in Ecology and Evolution. 27, 260–264 (2012).

14. Rubenstein, DR. *Animal Behavior*. New York, NY, Oxford University Press (2023).

15. Gowaty, PA. Field studies of parental care in birds: new data focus questions on variation among females. Adv. Study Behav. 25, 477–531 (1996).

16. Zahavi, A. Mate selection – a selection for a handicap. J. Theo. Biol. 53, 205–214 (1975).

17. Birkhead, TR, & Møller, A. Sperm Competition in Birds: Evolutionary Causes and Consequences. London, UK, Academic Press (1992).

18. Hamilton, WD, & Zuk, M. Heritable true fitness and bright birds: A role for parasites? Science. 218, 384–387 (1982).

19. Östlund, S & Ahnesjö, I. Female fifteen-spined sticklebacks prefer better fathers. Animal Behaviour. 56, 1177–1183 (1998).

20. Hill, GE. Plumage coloration is a sexually selected indicator of male quality. Nature. 350, 337–339 (1991).

21. Lewis, SM, & Cratsley, CK. Flash signal evolution, mate choice, and predation in fireflies. Annual Review of Entomology. 53, 293–321 (2008).

22. Patricelli, GL, Uy, JAC, & Borgia, G. Multiple male traits interact: Attractive bower decorations facilitate attractive behavioral displays in satin bowerbirds. Proc. R. Soc. B. 270, 2389–2395 (2003).

23. Kodric-Brown, A & Brown, JH. Truth in advertising: The kinds of traits favored by sexual selection. Am. Nat. 124, 309–323 (1984).

24. Endler, JA. *Natural Selection in the Wild*. Princeton, NJ, Princeton University Press (1986).

25. Morand Ferron, J, Cole, EF, & Quinn, JL. Studying the evolutionary ecology of cognition in the wild: a review of practical and conceptual challenges. Biological Reviews, 91, 367–389 (2016).

26. Shettleworth, SJ. Cognition, Evolution, and Behavior. New York, NY, Oxford University Press (2010).

27. Johnston, TD. Selective costs and benefits in the evolution of learning. In Advances in the Study of Behavior. Academic Press 12, 65-106 (1982).

28. Boogert, NJ, Fawcett, TW, Lefebvre, L. Mate choice for cognitive traits: a review of the evidence in nonhuman vertebrates, Behav. Ecol. 22, 447–459 (2011).

29. Snowberg, LK, & Benkman, CW. Mate choice based on a key ecological performance trait. J. Evol. Biol. 22, 762–769 (2009).

30. Chen, J, Zou, Y, Sun, Y-H, & Ten Cate, C. Problem-solving males become more attractive to female budgerigars. Science 363, 166 –167 (2019).

31. Shohet, AJ, & Watt, PJ. Female guppies *Poecilia reticulata* prefer males that can learn fast. J Fish Biol. 75, 1123 – 1523 (2009).

32. Spritzer, MD, Meikle, DB, & Solomon, NG. Female choice based on male spatial ability and aggressiveness among meadow voles. Anim Behav. 69, 1121–1130 (2005).

33. Shaw, RC, MacKinlay RD, Clayton, NS, & Burns, KC. Memory performance influences male reproductive success in a wild bird. Curr. Biol. 29, 1498–1502 (2019).

34. Keagy, J. Savard, JF, & Borgia, G. Male satin bowerbird problem-solving ability predicts mating success. Anim. Behav. 78, 809–817 (2009).

35. Croston, R, Branch, CL, Kozlovsky, DY, Dukas, R, & Pravosudov, VV. Heritability and the evolution of cognitive traits. Behav. Ecol. 26, 1447–1459 (2015).

36. McCallum, D, Grundel, R, Dahlsten, DL. Mountain Chickadee (*Poecile gambeli*), version 1.0. In Birds of the World (A. F. Poole and F. B. Gill, Editors). Cornell Lab of Ornithology, Ithaca, NY, USA (2020).

37. Pravosudov, VV, & Roth II, TC. Cognitive ecology of food hoarding: the evolution of spatial memory and the hippocampus. Ann. Rev. Ecol. Evol. System. 44, 173–193 (2013).

38. Lemmon D, Withiam ML, Barkan CPL. Mate protection and winter pair-bonds in black-capped chickadees. Condor 99, 424–433 (1997).

39. Ekman, J. Ecology of non-breeding social systems of Parus. Wil. Bull. 101, 263–288 (1989).

40. Sonnenberg, BR, Branch, CL, Pitera, AM, Bridge, E, & Pravosudov, VV. Natural selection and spatial cognition in wild food-caching mountain chickadees. Curr. Biol. 29, 670–676 (2019).

41. Welklin, JF et al. Spatial cognitive ability is associated with longevity in a food-caching bird. Science, 385, 1111–1115 (2024).

42. Branch, CL, Pitera, AM, Kozlovsky, DY, Bridge, ES, & Pravosudov, VV. Smart is the new sexy: female mountain chickadees increase reproductive investment when mated to males with better spatial cognition. Ecol. Lett. 22, 897–903 (2019).

43. Branch, CL, et al. What’s in a mate? Social pairing decisions and cognitive performance in food-caching mountain chickadees. Proc. R. Soc. B. 290, 20231073 (2023).

44. Otter, K, Ratcliffe, L, & Boag P. Extra-pair paternity in the black-capped chickadee. Condor 96, 218 – 222 (1994).

45. Mennill DJ, Ramsay SM, Boag PT, & Ratcliffe LM. Patterns of extrapair mating in relation to male dominance status and female nest placement in black-capped chickadees. Behavioral Ecology, 15, 757–765. (2004).

46. Heinen, VK et al. Social dominance has limited effects on spatial cognition in a wild food-caching birds. Proc. R. Soc. B. 288, 20211784 (2021).

47. Branch, CL et al. The genetic basis of spatial cognitive variation in a food-caching bird. Curr. Biol. 32, 210–219 (2022).

48. Semenov, G et al. Genes and gene networks underlying spatial cognition in food-caching chickadees. Curr. Biol. 34, 1930–1939 (2024).

49. Heinen, VK et al. Specialized spatial cognition is associated with reduced cognitive senescence in a food-caching bird. Proc. R. Soc. B. 288, 20203180 (2021).

50. Herdtle A, Duncan C, Manser MB, & Clutton-Brock T. Patterns and drivers of female extra-pair mating in wild Kalahari meerkats. Behavioral Ecology, 36: 3 (2025).

51. Neff BD, & Pitcher TE. Genetic quality and sexual selection: an integrated framework for good genes and compatible genes. Mol Ecol. 14, 19–38. (2005).

52. Bonderud ES, Otter KA, Burg TM, Marini KLD, Reudink MW. Patterns of extra-pair paternity in mountain chickadees. Ethology. 124, 378–386. (2018).

53. Hsu YH, Schroeder J, Winney I, Burke T & Nakagawa S. Are extra pair males different from cuckolded males? A case study and a meta analytic examination. Mol. Ecol., 24, 1558–1571 (2015).

54. Kozlovsky, DY, Branch, CL, Pitera, AM, Croston, R, & Pravosudov, VV. Fluctuations in annual climatic extremes are associated with reproductive variation in resident mountain chickadees. R Soc. Open Sci. 5, 171604 (2018).

55. Whitenack, LE, et al. Complex relationships between climate and reproduction in a resident montane bird. R. Soc. Open Sci. 10, 230554 (2023).

56. Croston, R et al. Individual variation in spatial memory performance in wild mountain chickadees from different elevations. Anim. Behav. 111, 225–234 (2016).

57. Tello-Ramos, MC et al. Memory in wild mountain chickadees from different elevations: comparing first-year birds with older survivors. Anim. Behav. 137, 149–160 (2018).

58. Benedict, LM et al. Information maintenance of food sources is associated with environment, spatial cognition and age in a food-caching bird. Anim. Behav. 182, 153–172 (2021).

59. Benedict, LM, Heinen, VK, Sonnenberg, BR, Bridge, ES, & Pravosudov, VV. Learning predictably alternating food locations across days in a food-caching bird. Anim. Behav. 196, 55–81 (2023).

60. Pyle, P. Identification Guide to North American Birds, Part I. Second Edition. Sheridan Books, Ann Arbor, MI, USA. (2022).

61. Thrasher, DJ, Butcher, BG, Campagna, L, Webster, MS, & Lovette, IJ. Double digest RAD sequencing outperforms microsatellite loci at assigning paternity and estimating relatedness: A proof of concept in a highly promiscuous bird. *Mol*. Ecol. Res. 18, 953–965 (2018).

62. Rohland, N, & Reich, D. Cost-effective, high-throughput DNA sequencing libraries for multiplexed target capture. Genome Res. 22, 939–46 (2012).

63. Wigginton, JE, Cutler, DJ, & Abecasis, GR. A note on exact tests of Hardy Weinberg equilibrium. Am. J Human Genetics 76, 887–893 (2005).

64. Danecek, P, et al. The variant call format and VCFtools. Bioinformatics 27, 2156–2158 (2011).

65. Gosselin, T. radiator: RADseq Data Exploration, Manipulation and Visualization using R. R package version 1.1.9 https://thierrygosselin.github.io/radiator/ (2020).

66. Kalinowski, S, Taper, M, & Marshall, T. Revising how the computer program CERVUS accommodates genotyping error increases success in paternity assignment. Mol Ecol. 16, 1099-1106 (2007).

67. Brooks, ME, et al. glmmTMB balances speed and flexibility among packages for zero-inflated generalized linear mixed modeling. R J. 9, 378–400 (2017).

68. Double, M, & Cockburn, A. Pre-dawn infidelity: females control extra-pair mating in superb fairy-wrens. Proc Biol Sci. 267, 465–70 (2000).

69. Joe, H, & Zhu, R. Generalized Poisson distribution: the property of mixture of Poisson and comparison with negative binomial distribution. Biometrical Journal: J. Math. Methods in Biosci. 47, 219–229 (2005).

70. Whitenack, L.E., Richmond, A., Sonnenberg, B.R., Welklin, J.F., Heinen, V.K., Pitera, A.M., Branch, C.L., Pravosudov, V.V. Postnatal dispersal and drivers of successful recruitment in resident Mountain Chickadees, Ornithology, ukaf006 (2025).

71. R Core Team. R: A language and environment for statistical computing (2021).

72. Hartig, F. DHARMa: Residual diagnostics for hierarchical (multi-level/mixed) regression models R package Version 0.4.4. (2018).

73. Welklin, JF et al. No evidence of reproductive senescence within natural life span in resident mountain chickadees (*Poecile gambeli*). Behav. Ecol. Sociobiol. 79, 10 (2025).

74. Smith, SM. Extra-pair copulations in black-capped chickadees: the role of the female. Behav. 107, 15–23. (1988).

75. Hasselquist, D, & Sherman, PW. Social mating systems and extrapair fertilizations in passerine birds. Behav. Ecol. 12, 457–466. (2001).

76. Kempenaers, B, Verheyen, GR, Vanderbroeck, M, Burke, T, Vanbroeckoven, C, & Dhondt, AA. Extra-pair paternity results from female preference for high-quality males in the blue tit. Nature 357, 494–96 (1992).

77. Sheldon, BC. Male phenotype, fertility, and the pursuit of extra-pair copulations by female birds. Proc. R. Soc. B. 257, 25–30 (1994).

78. Branch, CL, Kozlovsky, DY, & Pravosudov, VV. Elevation related variation in aggressive response to mirror image in mountain chickadees. Behav. 152, 667–676 (2015).

79. Kozlovsky, DY, Branch, CL, Freas, CA, & Pravosudov, VV. Elevation-related differences in exploration and social dominance in mountain chickadees (*Poecile gambeli*). Behav. Ecol. Sociobiol. 68, 1871–1881 (2014).

80. Branch, CL, et al. Testing the greater male variability phenomenon: male mountain chickadees exhibit larger variation in reversal learning performance compared to females. Proc. R. Soc. B. 287, 20200895 (2020).

81. Robayo Noguera, L, Stevenson, CAL, Wang, T, Pasquale, MK, Branch, CL. Variation in plumage reflectance but not song reflects spatial cognitive performance in black-capped chickadees *(Poecile atricapillus)*. Anim. Cog. 28, 14 (2025).

82. Haley SM, Sonnenberg BR, Heinen VK, Bridge ES, Pravosudov VV, Branch CL. More smarts, more song: male chickadees with better spatial learning and memory abilities sing more throughout the day. Proc. R. Soc. B. 293, 20260376 (2026).

83. Lane E, Kent KD, Agarwal M, Pravosudov VV, Branch CL. Is it a signal? Plumage reflectance is associated with winter climatic harshness and spatial cognition in a food-caching bird. Behav. Ecol. Sociobiol. In prep.

84. Lifjeld JT, Kleven O, Amundsen T, & Slagsvold T. Extrapair copulations in coal tits (*Periparus ater*): Female insurance against male fertility. Beh. Ecol. Sociobiol. 79, 79 (2025).

85. Canal D, Jovani R, & Potti J. Male decisions or female accessibility? Spatiotemporal patterns of extra pair paternity in a songbird. Behavioral Ecology, 23, 1146–1153 (2012).

86. Ratcliffe, L, Mennill, DJ, & Schubert, KA. Social dominance and fitness in black-capped chickadees in Ecology and Behaviour of Chickadees and Titmice: An integrative approach. Oxford, UK, Oxford University Press (2007).

87. Brodbeck, DR, Shettleworth, SJ. Matching location and color of a compound stimulus: comparison of a food-storing and nonstoring bird species. J. Exp.Psychol. 21, 64–77 (1995).

88. Pritchard JK, Pickrell JK, & Coop G. The genetics of human adaptation: hard sweeps, soft sweeps, and polygenic adaptation. Curr. Biol. 23, R208–R215 (2010).

89. Freas, CA, LaDage LD, Roth II, TC, & Pravosudov, VV. Elevation-related differences in memory and the hippocampus in mountain chickadees, *Poecile gambeli*. Anim. Behav. 84, 121–127 (2012).

90. Branch, CL, Jahner, JP, Kozlovsky, DY, Parchman, TL, & Pravosudov, VV. Absence of population structure across elevational gradients despite large phenotypic variation in mountain chickadees *(Poecile gambeli)*. R. Soc. Open Sci. 4, 170057 (2017).

91. Sonnenberg, B.R., Heinen, V.K., Pitera, A.M., Benedict, L.M., Branch, C.L., Bridge, E.S., Ouyang, J., Pravosudov, V.V. Natural variation in developmental condition has limited effect on spatial cognition in a wild food-caching bird. Proceedings of the Royal Society B 289, 1984 (2022).

92. Pitera, AM, Branch, CL, Sonnenberg, BR, Benedict, LM, Kozlovsky, DY, Pravosudov, VV. Reproduction is affected by individual breeding experience but not pair longevity in a socially monogamous bird. Behav. Ecol. Sociobiol. 75, 101 (2021).

93. Sonnenberg, BR, et al. Long-term winter food supplementation in a montane environment shows no significant impact on reproductive success in a resident food-caching bird. Ornithology 140, 1–12 (2022).

94. Branch, CL et al., Code and data for: Seeking smarts: Male chickadees with better spatial cognition sire more extra-pair young. Figshare (2026); 10.6084/m9.figshare.25343779

